# Klumpy: A Tool to Evaluate the Integrity of Long-Read Genome Assemblies and Illusive Sequence Motifs

**DOI:** 10.1101/2024.02.14.580330

**Authors:** Giovanni Madrigal, Bushra Fazal Minhas, Julian Catchen

## Abstract

The improvement and decreasing costs of third-generation sequencing technologies has widened the scope of biological questions researchers can address with de novo genome assemblies. With the increasing number of reference genomes, validating their integrity with minimal overhead is vital for establishing confident results in their applications. Here, we present Klumpy, a tool for detecting and visualizing both misassembled regions in a genome assembly and genetic elements (e.g., genes, promotors, or transposable elements) of interest in a set of sequences. By leveraging the initial raw reads in combination with their respective genome assembly, we illustrate Klumpy’s utility by investigating antifreeze glycoprotein (afgp) loci across two icefishes, by searching for a reported absent gene in the northern snakehead fish, and by scanning the reference genomes of a mudskipper and bumblebee for misassembled regions. In the two former cases, we were able to provide support for the noncanonical placement of an afgp locus in the icefishes and locate the missing snakehead gene. Furthermore, our genome scans were able to identify an cryptic locus in the mudskipper reference genome, and identify a putative repetitive element shared amongst several species of bees.

## Introduction

Since the publication of the first genome, the ϕX bacteriophage (Sanger et al. 1977), the number of publicly available genome assemblies has increased to over a million in 2021 (Sayers et al. 2021). A prominent feature accompanying the rate of assembly production was the three generations of sequencing technology underlying it. Two major transitions occurred, from massively-scaled, consortium-based, long-read assemblies, to an order of magnitude more independent productions of short-read, low quality assemblies, and most recently a transition enabling the production of very long-read assemblies by independent groups (Dijk et al. 2023). The benefits of long reads generated by Pacific Biosciences (PacBio) or Oxford Nanopore Technologies (ONT) continue to accrue in their ability to reveal more complex genome architectures (Marx 2023; Wohlers et al. 2023), with regions that have been notoriously difficult to assemble (e.g., long stretches of repeats, duplicate genes) becoming resolvable (Huddleston et al. 2014; Berná et al. 2018; Liu et al. 2019; Nath et al. 2021). Despite the advantages afforded by long reads, genome assembly remains a challenging undertaking due to genomic complexities such as ploidy, heterozygosity, and mosaicism (Jayakumar and Sakakibara 2019; Amarasinghe et al. 2020; Pucker et al. 2022; Hotaling et al. 2023).

Earlier iterations of long-read technologies generated reads with a relatively high level of insertion-deletion (indel) errors; to circumvent this issue, higher-accuracy short-reads were used to polish long-read contigs (Baptista and Kissinger 2019). However, with higher accuracy long-reads being generated, including PacBio HiFi sequencing (Hotaling et al. 2023), assembling a genome solely using long-reads has become routine. Inherently, there is a demand for reliable metrics to assess the quality of a genome assembly (Wang and Wang 2023). Such measures include the median contig or scaffold length (i.e., the N50 metric), gene space completeness through tools such as BUSCO (Benchmarking Universal Single-Copy Orthlog) and compleasm (Parra et al. 2009; Simão et al. 2015; Huang and Li 2023)., and in some instances, the detection and removal of exogenous contamination (Koutsovoulos et al. 2016; Wang et al. 2020; Sontowski et al. 2022; Diesel et al. 2023; Gao et al. 2023). For example, Mathers (2020) generated an improved assembly for the soybean aphid (*Aphis glycines* Matsumura) after the removal of parasitoid wasp contamination, while Wagner et al. (2023) used a combination BLAST+ tools (Altschul et al. 1990;Camacho et al. 2009) to remove non-Chondrichthyes and mitochondrial associated scaffolds when assembling the spiny dogfish (*Squalus acanthias*) nuclear genome. It is important for researchers to understand the factors that impact these values, as it has been reported that different genome assembly metrics are not necessarily correlated (Jauhal and Newcomb 2021) and in some cases, misleading (Bradnam et al. 2013; Rayamajhi et al. 2022). For instance, Rayamajhi et al. (2022) found inconsistency in both the gene duplications reported by BUSCO when varying the type of sequencing technology used for genome assembly, and the number of genes reported to be complete when changing the orientation of the genomic sequences.

Following assembly, genes are typically annotated. Caution must be taken when annotating a genome using gene models from related-species or inter- or intra-specific RNA-sequencing data. If the genome assembly itself contains errors in genic regions, these errors may cause adverse effects in the annotation process (Tørresen et al. 2019). Additionally, annotation pipelines may have difficulty with specific genes when their origin or structure is complex (Rust et al. 2002; Ejigu and Jung 2020; Mathé and Dunand 2021). For example, in Antarctic notothenioid fishes, antifreeze glycoproteins (*afgp*) prevent the growth of ice crystals *in vivo* and subsequently protect the fish from freezing in the Southern Ocean. However, the origin of the *afgp* genes lie in a set of Thr-Ala-Ala (or Thr-Pro-Ala) amino acid repeats of various lengths while the genes themselves occur as an array of tandem duplications (Chen et al. 1997a, 1997b; Kim et al. 2019; Bista et al. 2023), making their boundaries difficult to annotate (Zhuang et al. 2012). Given that the gene content of a *de novo* assembly is generally not known, gene annotation failures may be false negatives – occurring due to the failure of the annotation algorithm, or they may be true negatives – a biological characteristic of the sequenced individual (Nowoshilow et al. 2018). For candidate genes, it is important to able to distinguish between the two cases. Given the complexity of genome assembly methods, there continues to be interest in the development of algorithms for detecting misassemblies through different sources of information such as read coverage (Zhu et al. 2015), improper read alignments (Hunt et al. 2013), discrepancies in k-mer distributions between a reference genome and long-reads (Dishuck et al. 2023), mate-pair information (Kelley and Salzberg 2010), and notable breakpoints based on the alignment of the focal assembly, the constituent raw reads, or the genome of a closely related species (Bao et al. 2018; Asalone et al. 2020; Lai et al. 2022; Zhang et al. 2023).

Here, we present Klumpy, a bioinformatic tool designed to detect genome misassemblies, misannotations, and incongruities in long-read-based genome assemblies and their constituent raw reads. Klumpy is a Python package that can A) scan through a genome assembly and provide users with a list of potentially misassembled regions, and B) annotate sequences of interest (e.g., an assembled genome or its underlying raw reads) given a query of interest. These two modes of operation can work synergistically to C) annotate an assembly and the constituent raw reads together, based on a supplied, specific query (defined as any nucleotide sequence including, e.g., genes, regulatory motifs, or transposable elements). For instance, researchers may wish to search for a query within an assembly using BLASTN (Altschul et al. 1990); however, other software is often used to visualize the results when not using the web-based version (e.g., Kablammo; Wintersinger and Wasmuth 2015). In a like manner, when visualizing alignments, the Integrative Genomics Viewer (IGV; Thorvaldsdóttir et al. 2013) is often employed, which allows for the incorporation of gene annotations, though this feature is limited to annotations from the reference genome. Klumpy provides users the benefits of both tools, in which one can visualize “klumps” in both a reference genome and the aligned sequences in a sequence alignment producing easy to view images in PDF, SVG, or PNG format. After describing the algorithms of Klumpy, to illustrate its utility, we apply Klumpy to four different case studies, two of which start with a query gene of interest, and two in which we sought to discover misassembled regions with no prior knowledge of the assembled genome.

## Results

### Verifying the assembly and annotation of *afgp* genes in icefishes

In one of our recent studies, we sought to investigate the genomic changes that occur after a secondary adaptation to temperate environments in notothenioid fishes by assembling and comparing the genomes of *Champsocephalus gunnari* and *C. esox* (Rivera-Colón et al. 2023). Both species belong to the family Channichthyidae, a group of icefishes that lack hemoglobin and red blood cells (Sidell and O’Brien 2006), a physiological loss that is predicted to restrain these fishes to relatively highly oxygenated, cold waters. *C. esox* is the only documented icefish species found outside the Southern Ocean (Stankovic et al. 2002; Kock 2005) and yet it still possesses *afgp* coding regions (Miya et al. 2016). Given the role of *afgp* in the success of notothenioids in adapting to Antarctic waters (Matschiner et al. 2011), characterizing the *afgp* loci of *C. esox*, and its sister species, *C. gunnari*, is a key objective to elucidate the icefish polar-to-temperate transition. The assemblies were constructed using PacBio continuous long-read technology and annotated using BRAKER2 (Brůna et al. 2021); however, BRAKER2 failed to annotate any *afgp* loci in our assemblies. Using BLASTN, we manually annotated the *afgp* loci using an ortholog obtained from Nicodemus-Johnson et al. (2011). During curation, we found an additional, novel noncanonical *afgp* locus in both assemblies (Nicodemus-Johnson et al. 2011; Kim et al. 2019). This finding implies either 1) a second, novel *afgp* locus upstream of the primary locus, or 2) a genome assembly artifact, in which the assembler failed to correctly collapse the primary locus. We searched for *afgp* klumps in both the *C. gunnari* and *C. esox* assembled scaffolds, as well as in the respective raw reads. We used Klumpy to visualize the raw reads aligned against the scaffolds (both annotated with *afgp* klumps) and inspected the results for congruency.

The ‘find_klumps’ subprogram located 13 and 15 *afgp* klumps in the assembled chromosome 3 of *C. gunnari* and *C. esox* genomes, respectively (Figs. S1, S2, S3, and S4). Moreover, in both genome assemblies, Klumpy was able to uncover partial exons and complete exons of *afgp* (Tables S2 and S3). When viewing the alignments for *C. esox* at the noncanonical locus, two gaps bounded one of the *afgp* genes (Fig. 1A), which implied a potential misassembly (a Klumpy genome scan flagged this region when we applied the ‘scan_alignment’ subprogram on the *C. esox* reference genome). However, the reads at this position were able to span across the 3’ most gap and be mapped on the adjacent contig. Given that there are no alignments at the gap locations, our genome scan algorithm would not be able to construct a group that is able to tile across the gaps. Since the raw reads are able to span across the 3’ most gap (Fig. 1A), we could manually remove this gap from the assembly, however, since the 5’ gap is more ambiguous (several, soft-masked reads span it), it was left in the assembly. In the case of *C. gunnari*, there were no gaps in the noncanonical locus, as the alignments displayed an undisrupted tiling across the region despite the haplotypic differences visible in the alignments (Fig 1B). These results enabled us to successfully characterize this novel *afgp* locus and confirm that it was not a result of assembly error.

**Figure 1:**
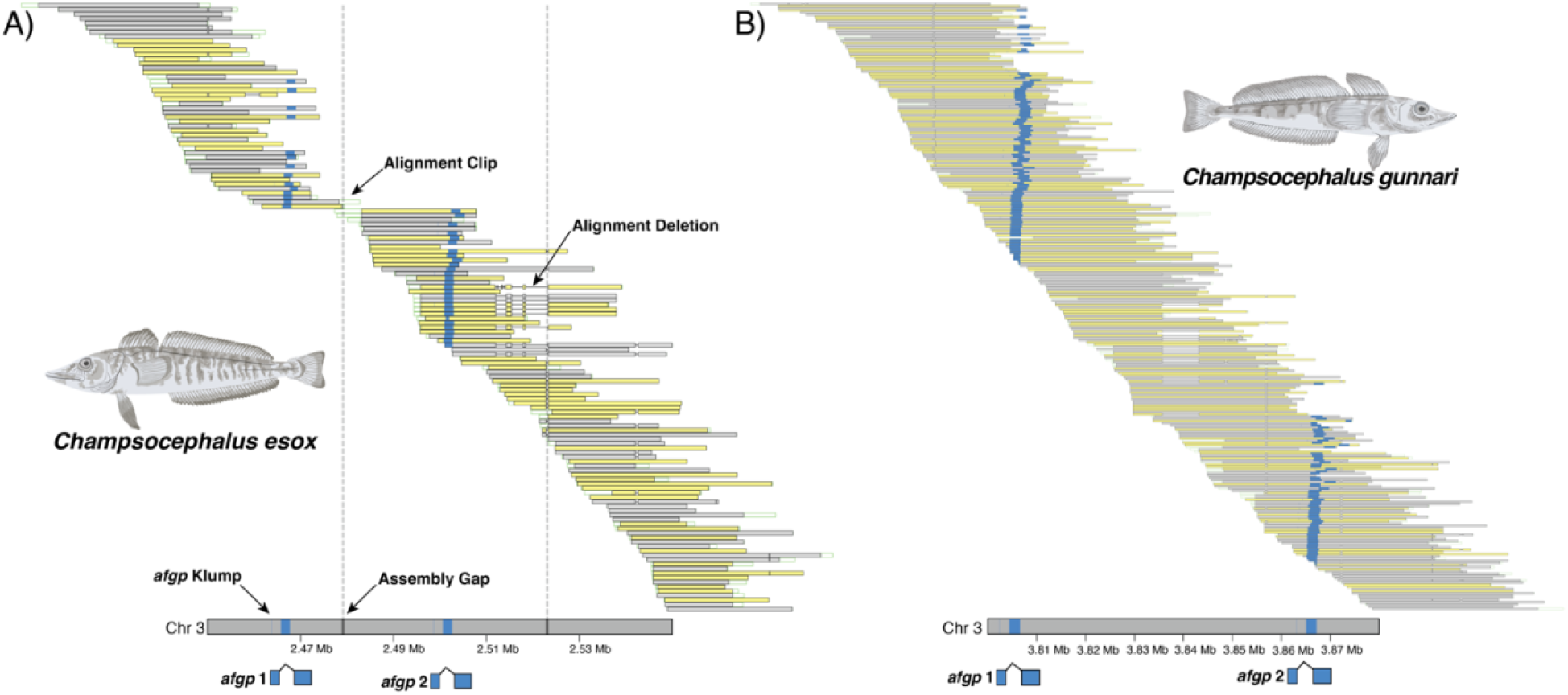
Alignment plot of the noncanonical locus for *C. esox* and *C. gunnari*. Yellow bars illustrate alignments to the reverse strand, and grey bars above the reference genome represent alignments to the forward strand. Blue boxes on the reference genome and alignments portray *afgp* klumps. A) vertical dash lines depict gaps in the assembly created during the scaffoldings. An *afgp* gene is contained in a single contig surrounded by two gaps, with reads mapping across the second and third contig confirming the joining of these two contigs. B) for *C. gunnari*, the alignments tile across the noncanonical locus, which presents strong evidence for this locus. In both cases, deletions (dash line connecting alignment blocks) are present in the some of the alignments, highlighting heterozygosity sites. For illustrative purposes, alignments shown are filtered in Klumpy by three criteria: a minimum sequence length of 15 Kbp for *C. gunnari* or 30 Kbp for *C. esox*, 75% of the sequence must be mapped.

We were further able to verify our manual annotation of the other *afgp* genes at the canonical sites (Figures S1 and S4).

### Elucidating the reported absence of the adcy5 gene in the northern snakehead

In our second example, we examined the reported absence of the adenylate cyclase 5 (*adcy5*) gene in a new, long-read-based assembly of the albino northern snakehead (*Channa argus*) genome (Zhou et al. 2022). The *adcy5* gene has been previously noted as having a functional role in guppy (*Poecilia reticulata*) pigmentation (Kottler et al. 2015), and its putative absence in the albino northern snakehead genome may reflect a similar effect on the organism’s phenotype. Interestingly, an alternative investigation by Sun et al. (2023) examined the molecular basis of the albino phenotype in the northern snakehead and proposed a non-sense mutation in the *slc45a2* gene as causal. We searched the results of Sun et al. (2023) and found no mention of *adcy5* or its association with the albino phenotype. Motivated by this discrepancy, we sought to apply Klumpy to search for the *adcy5* gene or any of its remnants in the albino northern snakehead reference genome to validate its association with the albino phenotype.

Using the exonic sequences of *adcy5* from the northern snakehead reference genome (a short-read assembly of the non-albino morph; Xu et al. 2017) as our query, we were able to locate the *adcy5* gene, consisting of 22 exons, on chromosome 10 of the albino, long-read assembly (Fig. 2). When we searched for a possible explanation to determine why the gene was reported absent by Zhou et al. (2022), we located an alternative codon, annotated in the albino reference but not in the non-albino reference, that contained a putative premature stop codon 528 bases upstream from the exon 1 klump (Fig. 2). Subjecting this annotated sequence to a BLASTN search, we identified this sequence as part of the *adcy5* gene, but it was missed in the short-read assembly. Interestingly, the premature stop codon and the surrounding base pairs are found in both northern snakehead assemblies (Fig. 2). We can speculate that the annotation was truncated in the albino genome because of the order the exons were encountered in the two genomes (the gene is annotated on alternative strands).

**Figure 2:**
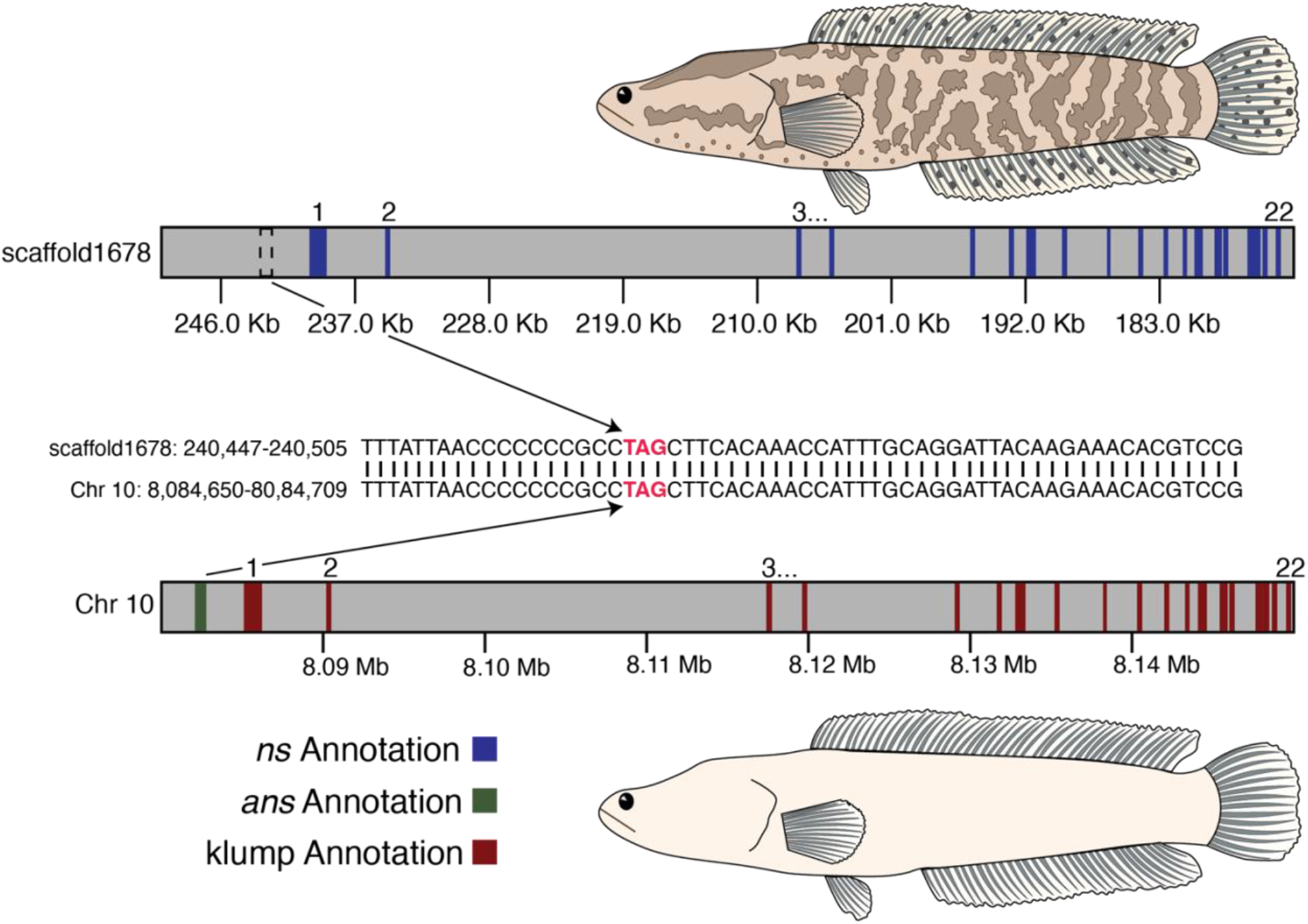
Depiction of the *adcy5* klumps and exon annotations in the wildtype (top) and albino (bottom) northern snakehead genome assemblies. The blue squares represent the annotated *adcy5* exons in the wildtype northern snakehead (*ns*) assembly, while the green square symbolizes the single annotated *adcy5* exon in the albino northern snakehead (*ans*) assembly. Maroon squares portray the *adcy5* exons identified by Klumpy. Numbers above the annotations correspond to annotations respective *adcy5* exon. Arrows stemming from the *ans* annotations point to a putative stop codon (red squares) that is found in both assemblies. Note that the *adcy5* gene is on the reverse strand in the Illumina assembly, while it is on the forward strand in the PacBio assembly.

In light of the alignment of the raw reads, we were unable to ascribe the mis-annotation of the *adcy5* gene to an assembly error on chromosome 10 (Fig. S5). The raw reads tiled across the entire *adcy5* gene and the nearest gap was located over 5.7 Mb away from the *adcy5* klumps. Furthermore, we were unable to replicate the syntenic region presented by Zhou et al. (2022) using Klumpy (Figs S6 and S7; Text S2). Regardless of these findings, a complete *adcy5* gene was located in the albino northern snakehead genome, supporting the findings by Sun et al. (2023).

### An anonymous locus in the great blue-spotted mudskipper genome

The great blue-spotted mudskipper (*Boleophthalmus pectinirostris*) is a species of interest for several research groups given its adaptations for inhabiting the terrestrial environment (Toba and Ishimatsu 2014; You et al. 2014, 2018; Storz et al. 2019; Kim et al. 2021). More generally, mudskippers have evolved morphological and physiological adaptations that allow for terrestrial locomotion (Hidayat et al. 2022), aerial vision and respiration (Sayer 2005), and immune defenses suited for terrestrial pathogens (You et al. 2014; Yi et al. 2017). Given these adaptations, mudskippers serve as a model for understanding the tetrapod water-to-land transition (Kutschera and Elliott 2013; Kim et al. 2021). In the study conducted by Bian et al. (2023), the authors generated three chromosome-level genome assemblies for mudskippers, including an improved reference genome of the great blue-spotted mudskipper. We applied Klumpy’s genome scan algorithm to examine the integrity of the assembly at coding regions and found a total of 3,119 regions flagged as incongruous between raw reads and assembled scaffolds. Klumpy examines the assembly by moving a window along assembled scaffolds, and interestingly, one window on scaffold NW_026571047.1 presented a case in which there was a lack of alignments within one of the introns of *insyn2a*. More specifically, there is a region containing cryptic sequence (i.e., a region with no coverage) where the alignments are soft clipped at the bounds of the locus (Fig. 3).

**Figure 3:**
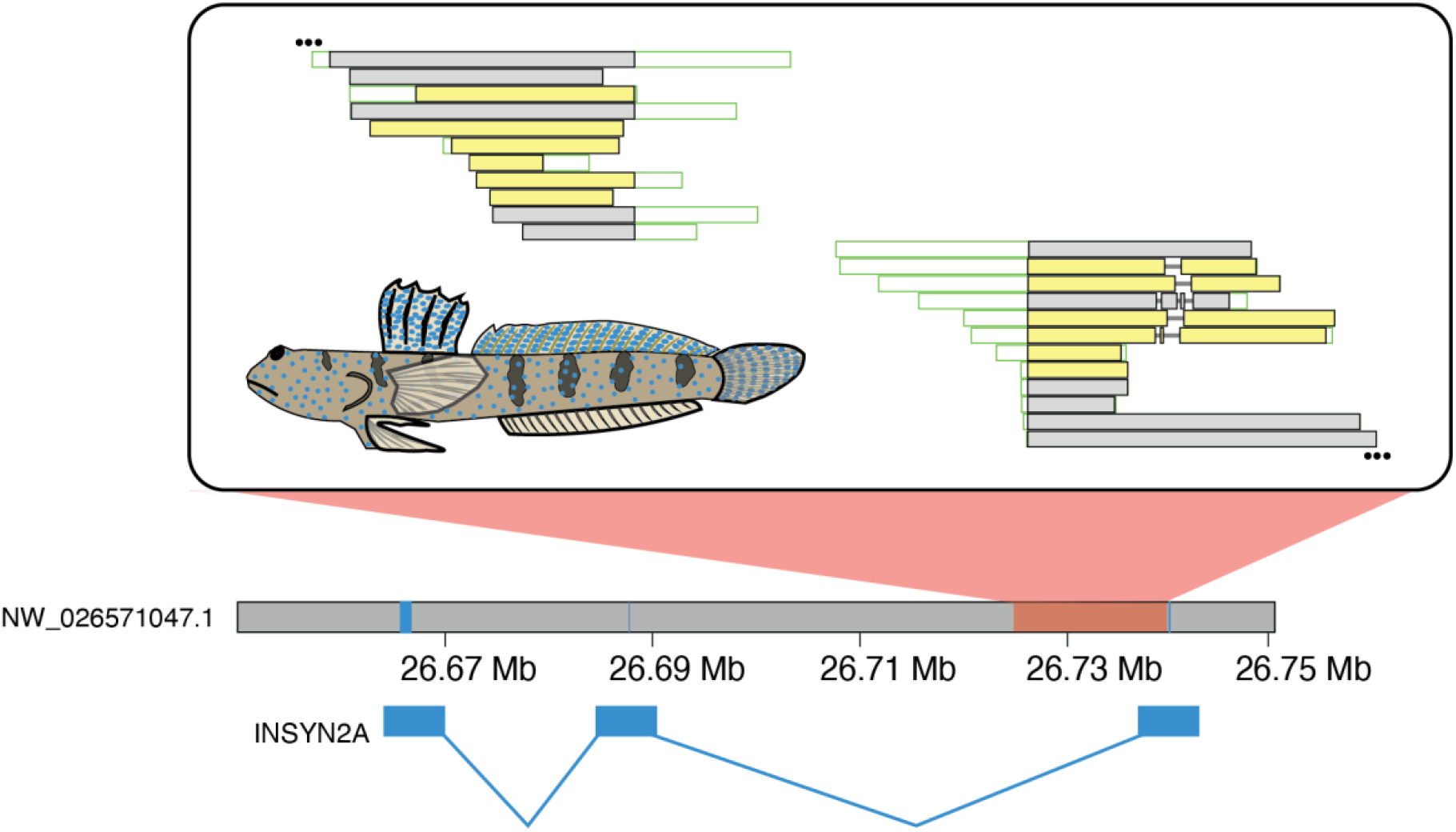
The *insyn2a* region containing a lack of coverage near the third exon in the great blue-spotted mudskipper genome assembly. Alignments flanking both sides of the unmapped region are clipped at the around the same position relative to the reference genome. Ellipsis at the top of the 5’ alignments and at the bottom of the 3’ alignments indicate a continuation of the alignments around *insyn2a*. A more comprehensive view of the alignments at this locus is presented in Fig. S11.

Centering on this region, re-alignment of the PacBio reads with Mummer 2.0 (Delcher et al. 2002) resulted in a similar outcome as the alignments generated by Minimap2 (Li 2018; Fig. S8), in which the reads could not be mapped across the locus. As the authors used short-read data to aid in the genome assembly process, we mapped the paired-end short-reads and found that this data set reflected the same alignment pattern found with the PacBio reads (Fig. S9). After performing a local assembly using the raw reads mapping between positions 26.65 Mb - 26.75 Mb on NW_026571047.1, a single contig containing the *insyn2a* gene lacking the cryptic sequence was constructed. Predictably, we could not map the contig back across the reference genome as it lacked the cryptic sequence (Fig. S10). Moreover, testing whether this cryptic sequence contained sequencing vector or contamination yielded no results, and BLASTN offered no additional insight to the origin of this sequence. Collectively, these results highlight a peculiar case in which a segment of the great blue-spotted mudskipper reference genome is untraceable whether using publicly available databases or the constituent raw reads and we putatively conclude that this is an artifact of the assembly process and not a true part of the genome.

### A problematic repeat in blumblebees

The Hunt’s bumblebee (*Bombus huntii*) has garnered a considerable amount of attention partially due to its pollinating capabilities (Xu et al. 2013; Strange 2015; Bobiwash et al. 2018; Koch et al. 2018; Baur et al. 2019) and as a resource for agricultural applications (Childers et al. 2021). We applied Klumpy’s genome scan to check the quality of the *B. huntti* reference genome and found 98 incongruous regions. We took a particular interest in a 75 Kbp region on chromosome 18 containing LOC126875579 (Fig. 4A). Using its exonic sequences, we found multiple klumps for each of the three exons across the *B. huntti* reference genome (Fig. S13) that were located near gaps in the assembly or at chromosomal end regions.

**Figure 4:**
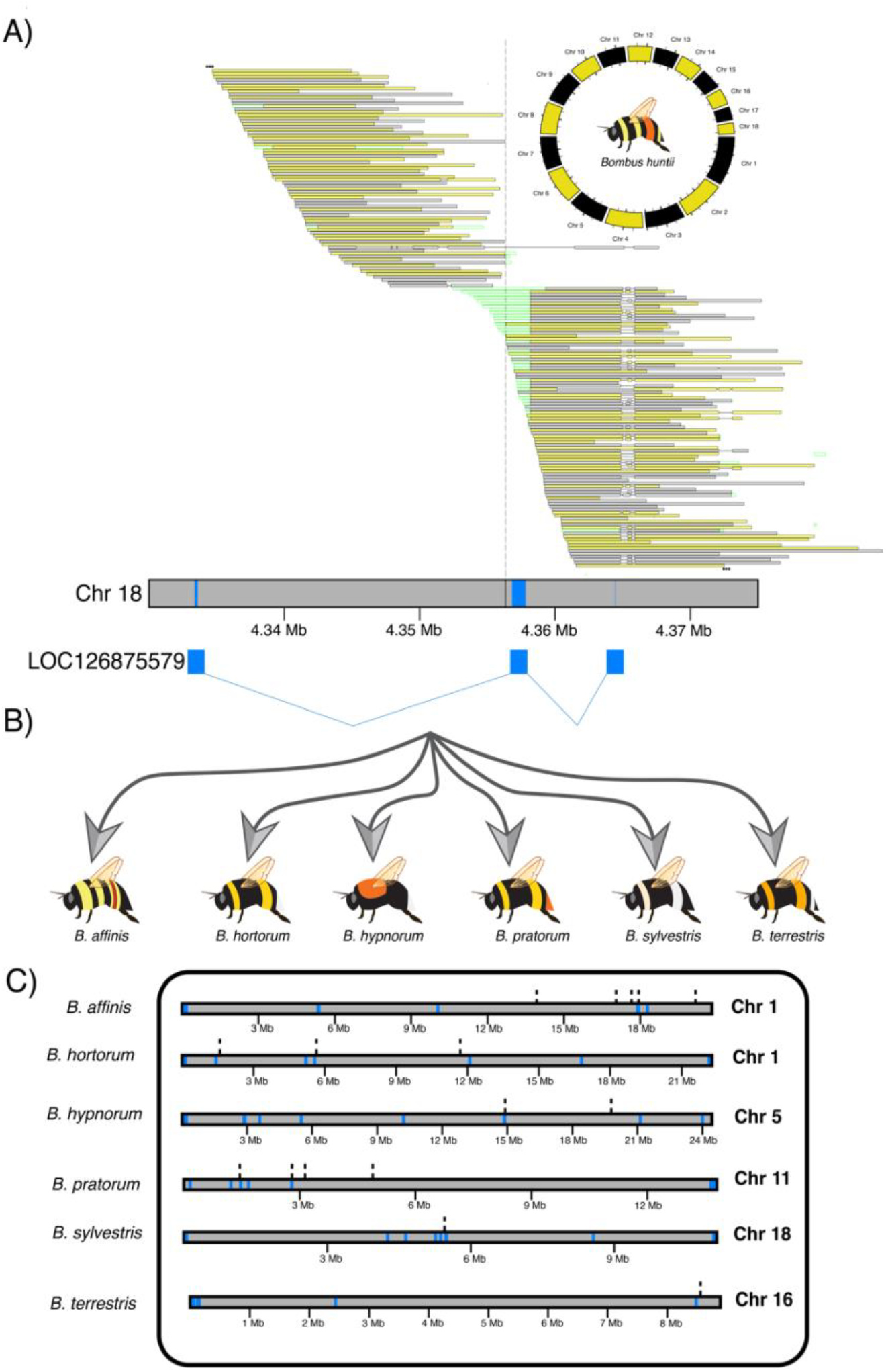
An overview of case study 4. A 75 Kbp window containing LOC126875579 was flagged when scanning the *Bombus huntii* reference genome for misassemblies. A) a gap is shown to split the three exons (light blue boxes) across two contigs. B) the exons were then used to locate LOC126875579 klumps across six other *Bombus* species. C) a similar pattern of LOC126875579 klumps (light blue boxes) across the *Bombus* assemblies, where the klumps can be found at the ends of the chromosomes and located nearby gaps (dashed lines). For illustration purposes, we only present one chromosome per species.

A BLASTN search of LOC126875579 did not yield any previously annotated orthologs. Interestingly, the BLASTN results were predominately composed of records from congenic species. Expanding our query search of LOC126875579 across six additional *Bombus* species (Fig. 4B), we found LOC126875579 klumps to be present in a similar manner as that found in *B. huntti*. Although the number of klumps varied across species (98 in *B. affinis*, 77 in *B. hortorum*, 419 in *B. huntti*, 306 in *B. hypnorum*, 435 in *B. pratorum*, 201 in *B. sylvestris*, and 109 in *B. terrestris*), the majority of the klumps were tandemly duplicated and several were near gaps or the ends of chromosomes in the assemblies. Additionally, with the exception of the *B. pratorum* and *B. terrestris* reference genomes, several unplaced scaffolds were shown to encompass tandemly duplicated LOC126875579 klumps (Figs. S13 - S19). Together, these results highlight a repetitive genetic element that is shared across multiple *Bombus* species that seems to locate itself within proximity of gaps and chromosome ends.

## Discussion

Improvements in long-read sequencing technology is driving commensurate improvements in reference genome assembly, however the need to properly assess assembly quality remains. Despite the breadth and complexity of genome assembly errors, most reported measures of assembly quality continue to be summary metrics such as N50 or summary reports of partial annotation results, such as BUSCO. Although several bioinformatic tools exist to detect misassembled regions (Meng et al. 2022), the complexity of these tools, with a reliance on multiple input files (Kelley and Salzberg 2010) or software dependencies (Zhang et al. 2023) may limit their use. Regardless, it is essential that assembly quality metrics successfully communicate the true status of an assembly. While metrics such as N50 and BUSCO may provide a qualitative sense of assembly quality, they do not signal to researchers specific problem areas and rarely the true magnitude in the number of problem areas.

In this study, we present Klumpy; a genome assembly quality assessment tool that may be incorporated natively into any third-generation genome assembly pipeline. Our intuitive approach of leveraging the tiling of alignments from the source data across the genome, alongside k-mer analysis and built-in visualization of these analyses, provides investigators a straightforward process to assess their genome of interest with minimal understanding of the organism’s molecular biology. In contrast, biologists with prior knowledge of the genomic characteristics associated with the focal taxa or related species may take advantage of homologous sequences from outside resources to assess known problematic loci (i.e., loci known to be difficult to assemble or annotate).

To illustrate Klumpy’s applicability, we demonstrated its usage in four separate case studies. In the first case, we were able to annotate a complex set of *afgp* gene copies that were missed by the automated annotation pipeline and verify that a new, novel *afgp* locus was real, and not an assembly artifact in the *C. gunnari* and *C. esox* (Fig. 2) assemblies. By annotating and viewing the aligned raw reads and assembled scaffolds together, we are able to deduce the reliability of the alignments and presence of the *afgp* sequences in the raw reads. In addition, the visualization of the alignments allowed us to understand when alignment indels were most likely to represent heterozygosity among the two chromosome copies (haplotigs), versus assembly error. Together, these results provided support for the *afgp* loci in our icefish genome assemblies and allowed us to report the first noncanonical *afgp* locus in the notothenioid literature (Rivera-Colón et al. 2023).

When employing Klumpy to survey the albino northern snakehead genome and the corresponding raw reads for the *adcy5* gene, Klumpy was able to confidently locate and annotate our gene of interest. Using these results, we were able to locate a putative premature stop codon upstream from the *adcy5* klumps, which may be the likely cause for the majority of the gene being unannotated. However, the sequence for the reported annotation of the partial *adcy5* locus in the albino northern snakehead genome assembly is also found in the phenotypically wildtype genome assembly of the northern snakehead. Given that the two genome assemblies were annotated using two different annotation pipelines and data, it is plausible that some genes may have been annotated in one assembly and left unannotated in the other. Another cause for the reported absence of the *adcy5* gene could be the specific gene identification methodology the authors applied, in which genome specific (i.e., non-albino vs albino phenotype) genes were determined by identifying genes with less than 80% overlap between coding sequences obtained from the two assemblies (Zhou et al. 2022). By using a shortened coding sequence of the *adcy5*, the *adcy5* gene would have been reported absent following the above approach. In cases like this, Klumpy will serve as an advantageous tool for verifying the absence of a gene.

To test our hypothesis-free, genome scan algorithm, we examined the great blue-spotted mudskipper PacBio assembly and found a locus containing cryptic sequence within one of the introns of the *insyn2a* gene. We scrutinized this locus using different methods but were unable to identify its source of origin. One explanation is the possibility of the locus being composed of fragments of multiple reads, which may have been generated as the assembler traversed an ambiguous junction in the assembly graph. This could be investigated reducing the k-mer size during the mapping phase to identify the corresponding segments from the raw reads (Noble and Pozhitkov 2018). Lastly, despite this sequence residing outside the exonic sites of *insyn2a*, its presence in the intronic sites may have consequential effects on downstream analyses of this gene. Ongoing research on introns have revealed their role in gene regulation (Rose 2019) and pre-mRNA splicing (Pagani and Baralle 2004), highlighting the importance for proper and accurate gene assemblage.

For our last case study, we explored the *Bombus huntti* genome for misassembled regions. In our search, we found the LOC126875579 exons to be on two separate contigs, which sparked a further investigation into this genomic feature. Upon expanding our analysis to include six more members of *Bombus*, we found the putative exons of LOC126875579 to be tandemly duplicated and nearby the gaps or ends in all the bumblebee genomes surveyed. Given the locations and organization of the LOC126875579 klumps, we predict LOC126875579 to be a clade-specific, transposable element as opposed to a protein-coding gene. As the BLASTN results are largely composed of sequences from the genus *Bombus*, we speculate that LOC126875579 originated from an ancestral *Bombus* species. It must be noted that several of the *Bombus* data sets used in this study have been documented to share a common 10-mer amongst other *Bombus* genome assemblies (Crowley et al. 2023b), with our results exemplifying a similar case. Despite these details, these chromosome-level genome assemblies remain useful and with the sequencing of additional bee species, the extent of which LOC126875579 can be found across *Bombus* and relatives can be revealed.

Despite the benefits Klumpy provides, several factors, both biological and technical, present appreciable obstacles to our algorithms. For instance, when haplotypic differences at a given locus are large enough, the alignments of the reads representing these haplotypes may not evenly tile across the focal site; particularly when the number of reads is not equally distributed between the different alleles. Additionally, the gaps and ends of chromosomes in a genome assembly may present a difficult region to properly align a sequence, which would inflate the number of reported windows in our genome scan algorithm. As the number of available genome assemblies greatly exceeds the number of genome assemblies we evaluated, it is likely that our genome scan algorithm also does not cover all cases of alignment patterns. Several of the reported windows in our case studies were discredited as false positives, indicating that our algorithm’s inclination towards false positives over false negatives favors a conservative approach. For example, the first 50 Kbp of scaffold NW_026099416.1 in the *B. huntii* reference genome was flagged due to a deletion in some of the alignments mapped to two small subunit ribosomal RNA loci; however, we expect this to not be a misassembly as there are reads at this site supporting both alternative alleles, but the amount of overlap between the consistent alignments (in respect to the reference genome) may have not been sufficient enough for Klumpy to consider this region correct (Fig. S20). Additionally, reads with many small indels called by the aligner will not be explicitly drawn (when the indel length is shorter than the --deletion_len parameter) and may lead to over-shifting the klump annotations when visualized in the ‘alignment_plot’ subprogram.

Another source of difficulty is the detection of small queries. Exons have been reported in numerous lengths, with the shortest reported exon composed of a single nucleotide (Guo and Liu 2015). Short exons present a challenge for annotation pipelines given their reduced specificity when compared to longer sequences (Mathé et al. 2002; Saeys et al. 2007; Mathé and Dunand 2021). In the case of the northern snakehead, the 17^th^ exon of *adcy5* was composed of 21 nucleotides, and thus to surmount this issue, we applied an additional Klumpy filter that required sequences to have klumps from multiple query sources (i.e., more than one exon) when surveying the genome assemblies. Similarly, sequences containing motifs that are common throughout the genome will yield a copious number of false positives. These false positives may result in partial matches; however, as in the case of the *afgp* loci in our analysis of icefishes, one must carefully examine these partial matches in order to avoid the removal of true positives. Despite these shortcomings, the inclusion of prior knowledge of the genomic characteristics of the organism or that of related taxa, may help guide researchers towards a more amenable analysis.

Altogether, Klumpy is an effective tool to assess the quality of a genome assembly and locate genes of interest. By combining the k-mer analysis properties of popular software such as BLASTN, with the visualization aspects of IGV, Klumpy gives users an efficient and desirable method for investigating genomic regions of interest. Equivalently, with Klumpy, one does not need to have prior knowledge of possible misassembled regions, which provides users with a starting point for evaluating a genome of interest. As we demonstrated with *Bombus huntti*, exploration of one genome may provoke further exploration of a gene or group of organisms, thus Klumpy is able to kindle scientific inquiries that are yet to be explored. Furthermore, by identifying misassemble regions, Klumpy offers users the opportunity to extract reads at such locations, which can then be locally reassembled into a primary contig using genome assemblers like Flye (Kolmogorov et al. 2019) or Hifiasm (Cheng et al. 2021) and re-incorporated back into the reference genome by applying a tool such as RagTag (Alonge et al. 2022). In combination with such tools, Klumpy will prove advantageous for assessing future and previous long-read genome assemblies.

## Materials and Methods

Klumpy has two major modes of operation depending on the user’s goals (Fig. 5). In the first mode, Klumpy scans through a set of long-read alignments to the contigs or scaffolds of a genome to identify regions where the raw reads are incongruent with the underlying assembly, indicating that the region may not be assembled correctly **-** flagging these regions. In the second mode of operation, Klumpy will search for a query sequence among a set of subject sequences by k-merizing the query and subjects. Subject sequences, in this case, may be the contigs or scaffolds of a genome assembly or a set of raw, long-reads. Clusters of shared k-mers – or klumps – are flagged. Klumpy can produce visualizations from either mode of operation and provides the user the ability to incorporate additional assembly features (i.e., assembly gaps and annotated genes) into the generated illustrations.

**Figure 5:**
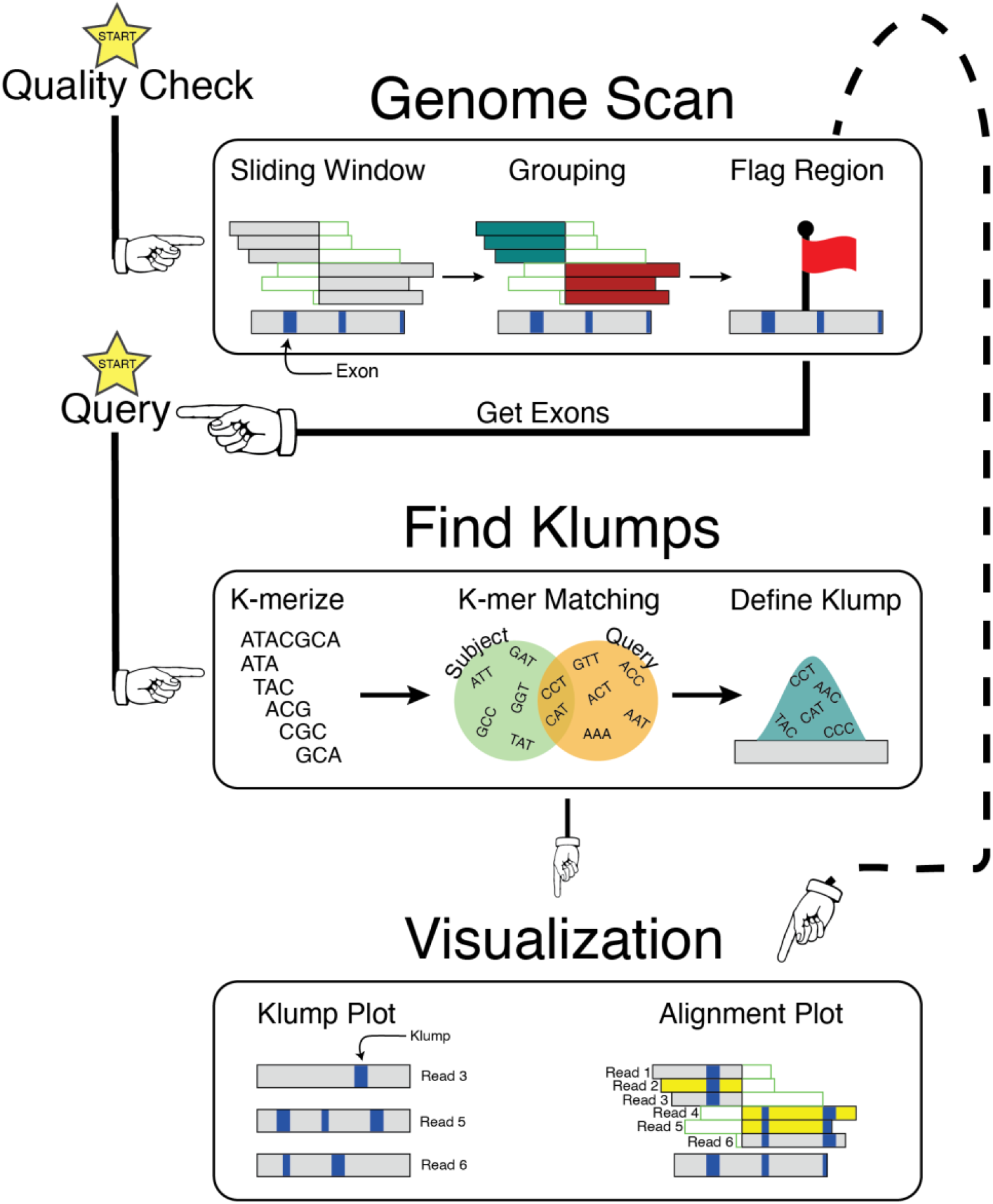
Synopsis of the Klumpy pipeline. There are two possible routes to start an analysis in Klumpy (yellow star icons). Starting with Quality Check, a sliding window approach is implemented to identify candidate misassembled regions. A grouping algorithm is performed on the alignments at each window, and in the case where no group can completely tile across the window, the window is flagged. If there are exons in the window (blue rectangles) that are of interest, exons can be retrieved from the assembly and used as a query. When executing the pipeline starting with a query, both the query and subject sequences will be k-merized, and k-mers shared between the two data sets will be used to locate klumps in the subject sequences. Klumps can then be viewed independently or used to annotate an alignment.

### Genome scanning mode of operation

To begin a genome scan, the user provides Klumpy with a set of alignments in SAM or BAM format (Li et al. 2009). Typically, these alignments will compare the raw reads submitted to the genome assembler to a *de novo* assembly of those reads. Klumpy will examine the genome in *windows*, where each window consists of a set of extracted alignments, ordered by their alignment positions in the genome. The initial size of the window is 50 Kbp or can be supplied by the user. The Klumpy grouping algorithm will be applied to each window (described below) and the window is then advanced along the scaffolds of the genome. The user may also supply the reference genome’s annotation file in GTF or GFF3 format that defines the gene locations, in which case, the window will advance to only encompass regions surrounding exonic sequences. During the grouping process, Klumpy will determine if the reads within a particular window are inconsistent and likely indicate an assembly anomaly.

For each window, Klumpy will extract primary alignments using Samtools (Li et al. 2009) that are within the window range. Reads are often clipped by an aligner and this clipping is described in the SAM file. Since clipped sites are not considered part of the alignment, Klumpy extends the targeted range in both directions so that read alignments possessing clipped bases at the edges of the window are incorporated. After setting the window size, the lengths of the assembly scaffolds are compared to that of the window size. Any reference contig/scaffold shorter than the window are removed from the analysis, which leaves the sequence alignments at those regions unprocessed. When filling a window, poor alignments – those that have a sequence length or percentage of aligned bases below the given thresholds – are excluded from the analysis. Similarly, the user can specify a minimum length and mapped percentage needed to include an alignment. Reads that are deemed untrustworthy remain in the data set but are excluded from the subsequent grouping process. Once the window is populated, if the number of reads is below a minimum threshold, the window is discarded. In a different manner, if the number of reads exceeds a maximum threshold and subsampling is permitted, the reads will be randomly subsampled to the maximum threshold. These windows tend to span repetitive regions, which are computationally expensive to process since each alignment is evaluated during the grouping process. In practice, subsampling has not negatively impacted the results as we anticipate there to be enough alignments to span the entire window if the region was correctly constructed. Once these checks have been performed, the data in the window is evaluated.

### The Klumpy grouping algorithm

Reads within a window are *grouped* according to their alignment patterns (Fig. 6) under the assumption that windows that contain more than one group demonstrate that more than one locus was collapsed during the assembly process. This may include heterozygous features in the underlying individual, such as insertions, deletions, or inversions, or it may include complex repeat sequences that were over-merged.

**Figure 6:**
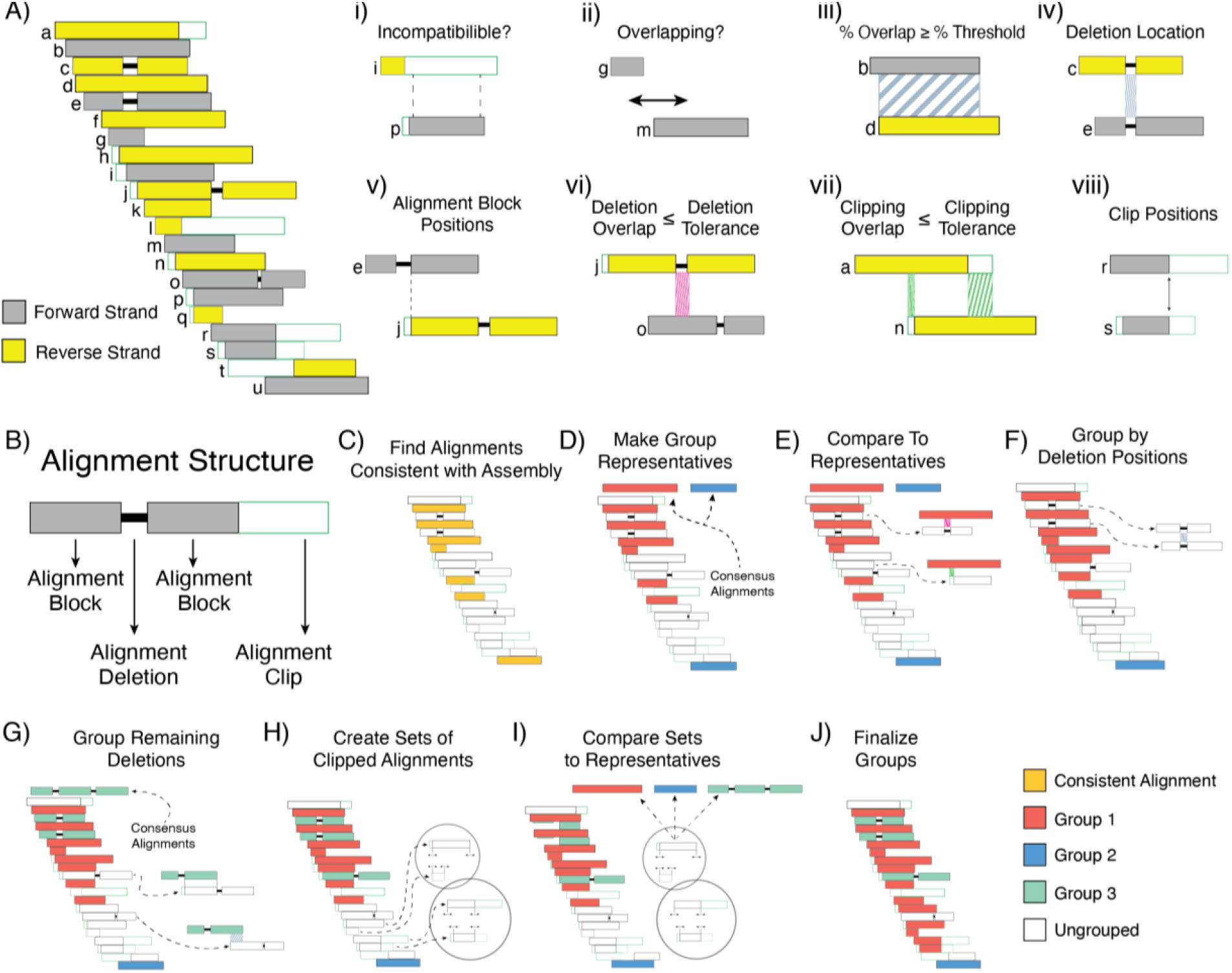
Alignment grouping algorithm. Given a set of alignments (A), each alignment’s structure (B) is examined and used to collect pieces of evidence through a pairwise comparison (i-viii) to predict which alignments are derived from the same locus. An initial check for incompatibility is performed by checking whether one of the alignments falls into the clipped portion of the other alignment (i). If incompatibility is not overt, whether there is any overlap in aligned positions between the two alignments (ii) dictates the continuation of this process. Evidence is collected by checking the percentages of overlap between the two alignments (iii), observing whether the alignments contain deletions at the same positions (iv), inspecting the alignment block positions (v), examining the tolerance thresholds for alignment deletions and clips between the two alignments (vi, vii), and surveying the positions of any clipped sites (viii). To construct the groups, consistent alignments (C) are used to initialize the first group(s) and a representative alignment for each group (D). The algorithm then attempts to add the ungrouped alignments by comparing their structures to the consensus alignment(s) (E). If there remain inconsistent alignments containing deletions at the same positions, these alignments are then used to propose one or more new groups (F). A pairwise comparison across all alignments possessing at least one deletion is then performed to establish the proposed groups, which leads to the formation of one or more consensus alignments (G). Next, sets of unincorporated clipped alignments are assembled by reviewing the sites where the alignments are clipped (H). The sets of clipped sequences are then compared to the present groups for possible incorporation (I), at which point the grouping algorithm is terminated after removing groups not containing the minimum number of alignments required to form a group (J). A list of the lines of evidence along with their weights is shown in Table S1.

Alignments to the assembled genome may be consistent or inconsistent – e.g., soft- or hard-clipped, or extended via deletions (Fig. 6B). The grouping algorithm begins by comparing reads pairwise and identifying sets of reads whose alignments are consistent with the assembled genome (Fig. 6C). These sets of reads are considered *groups* and consensus sequences are generated to represent each group (Fig. 6D). Klumpy then compares the remaining reads against each of these groups (Fig. 6E) and incorporates reads that may have small differences with the assembled genome but are still consistent with the group. This is done by considering each of the factors shown in Fig 2i-viii, which are explained in detail in Text S1 and Table S1. Next, sets of reads that share common deletions are grouped (Fig. 6F) and these deletion groups are merged if compatible with one another (Fig. 6G). Finally, sets of reads that contain large, clipped regions are compared against each other and merged when consistent. If a group contains too few reads, it is discarded. In cases where adjacent windows are flagged, windows will be collapsed into a single record.

The final set of groups in a window describes the integrity of a region of the assembly. One group in one window demonstrates consistency between the reads at that location. However, two, disconnected groups illustrate a region of the assembly that is not supported by raw reads. Two or more groups may also describe more complex features, such as an insertion, that have been collapsed in the assembly. The user can specify the number of groups (default 2), and Klumpy will flag and output windows that contain groups equal to or are above the threshold where no group is able to tile across the region. These groups can subsequently be visualized by the user in Klumpy, or the raw reads can be exported for further investigation (e.g., in order to assemble them independently and further examine the region).

### Generating query klumps

To find queried sequences (e.g., gene sequences) in a set of subject sequences (e.g., a reference genome or a collection of raw reads), Klumpy starts by compiling every unique subsequence of a size *k* (i.e., a k-mer) for each query sequence. Any queried sequence that is shorter in length than *k* is not incorporated into the subsequent analysis. The query k-mers are then searched for, identified, and located in the subject sequences. Once all the queried k-mers have been mapped onto the subject sequences, a klump can be defined by setting the minimum number of query k-mers that are no further apart than the set maximum distance in base pairs (bp). K-mers originating from a different source query are not included within an individual klump; albeit multiple klumps from different query sources at the same locus are permissible. The results are reported in a tabular formatted file that can be viewed by the user or resupplied to Klumpy for visualization.

### Investigation of *afgp* genes in icefishes

To determine the integrity of the *afgp* loci in the reference genomes of *C. gunnnari* and *C. esox*, we employed the ‘find_klumps’ subprogram of Klumpy, supplying the *afgp* model from Nicodemus-Johnson et al. (2011) as the query (--query) and mapping the *afgp* k-mers onto the genome assemblies (--subject). The analysis was performed individually for each species and repeated using the constituent raw reads as the subject sequences. We subsequently set the minimum number of k-mers (--min_kmers) per klump to 10 and the maximum range (--range) between klumps as 1 Kbp. The raw reads were aligned to their respective genome assembly using minimap2 (Li 2018) and processed with Samtools (Li et al. 2009) to create sorted BAM files. Additionally, gaps were located in the reference genome using the Klumpy subprogram ‘find_gaps’ using the --fasta option. These results were then visualized using the ‘alignment_plot’ Klumpy subprogram.

### Searching for the adenylate cyclase 5 (*adcy5*) gene in the northern snakehead

To obtain the *adcy5a* gene model, we extracted the exons from the northern snakehead reference genome (Xu et al. 2017) via the ‘get_exons’ Klumpy subprogram by supplying the program with the reference genome (--fasta), the annotations (--annotation) and the desired gene name (--genes). We then downloaded the raw reads from the NCBI Short Read Archive (accession: SRR8607404) and obtained the albino northern snakehead reference genome (accession GCA_004786185.1) from NCBI. *Adcy5* klumps were then searched for in both the reference genome and raw reads by applying the Klumpy subprogram ‘find_klumps’. The maximum distance between klumps was set to 50 bp (--range) and for klumps to be composed of at least 3 k-mers (--min_kmers). When analyzing the reference genome, we applied an additional filter to remove sequences that did not contain k-mers from at least 5 exons (--query_count) to exclude sequences containing spurious query matches.

To determine if the candidate *adcy5* region may be misassembled, an alignment of the raw reads to the reference genome was performed using minimap2 (Li 2018). The alignments were then sorted and indexed using Samtools (Li et al. 2009). Additionally, we used MUSCLE (Edgar 2004) to perform a local alignment between the two northern snakehead reference genomes upstream of the exon 1 klump to identify mechanisms that may have attributed to the findings of Zhou et al. (2022). Lastly, to reproduce the findings of Zhou et al. (2022), we repeated the ‘find_klumps’ analysis with the Illumina-based northern snakehead assembly.

### A genomic scan on the great blue-spotted mudskipper genome

We downloaded the great blue-spotted mudskipper PacBio assembly (GCF_026225935.1) and used the SRA-toolkit to obtain the raw PacBio reads (SRA accession: SRR20746324). The alignments of the raw reads were generated using minimap2 (Li 2018) using the same approach as performed with the northern snakehead case study. We next performed the genome scan using the subprogram ‘scan_alignment’, using default values, with the exceptions of supplying the software with the reference genome annotations so that only windows containing exonic sequences would be evaluated.

The reported windows were then inspected to validate our method. We focused on a region in scaffold NW_026571047.1 containing the inhibitory synaptic factor 2A (*insyn2a*) gene to elucidate the source from which the locus was constructed. The short-read data used for this assembly (SRA accessions: SRR20746323, SRR20746325, SRR20746326, SRR20746327) were mapped to the mudskipper genome using BWA mem (Li 2013) and the region was again inspected using the subprogram ‘alignment_plot’. Furthermore, we aligned the PacBio data to the unmapped locus using Mummer2 (Delcher et al. 2002) to determine the influence of the alignment algorithm on our finding. We then performed a local reassembly using Flye (Kolmogorov et al. 2019) with the reads that mapped between 26.65 Mb - 26.75 Mb on NW_026571047.1 as an attempt to recreate the focal locus. The resulting contig was then mapped back to the reference genome using minimap2 (Li 2018). To establish whether the region was a result of sequence contamination, the locus was extracted from the reference genome and processed through the VecScreen pipeline (https://www.ncbi.nlm.nih.gov/tools/vecscreen/). Additionally, we subjected these same nucleotides to a web-based BLASTN search to identify homologous loci.

### An exploration into bumblebee genome assemblies

In the last case study, we explored the quality of the Hunt’s bumblebee reference genome (Childers et al. 2021). The reference assembly was obtained from NCBI (accession GCF_024542735.1) and the SRA-toolkit was employed to retrieve its corresponding PacBio reads (SRA accession: SRR20215638). Minimap2 (Li 2018) was used to map the raw reads to the reference genome and the alignments were processed through Samtools (Li et al. 2009) to sort and index the records. We then replicated the same parameters used in our mudskipper analysis to scan the reference genome.

Upon exploration of the reported windows, we found LOC126875579 of particular interest. We extracted the exonic sequences of LOC126875579 using ‘get_exons’ by supplying Klumpy with the reference genome and annotations. The exons were then searched for to identify similar sequences in the reference genome using the ‘find_klumps’ Klumpy subprogram. More specifically, we formed klumps consisting of at least 11 k-mers with the maximum distance cutoff between klumps as 50 bp. To identify orthologs, we used BLASTN to search the NCBI NR database using the LOC126875579 exons as the query. In light of the BLASTN results, we searched the genome assemblies of *B. affinis* (GCF_024516045.1; Koch et al. 2023), *B. hortorum* (GCA_905332935.1; Crowley 2021), *B. hypnorum* (GCA_911387925.2; Crowley et al. 2023a), *B. pratorum* (GCA_930367275.1; Crowley et al. 2023c), *B. sylvestris* (GCA_911622165.2; Crowley 2023), and *B. terrestris* (GCF_910591885.1; Crowley et al. 2023b) for LOC126875579 klumps. To keep our analysis consistent across all bee species, we generated klumps using the same parameters applied to *B. huntti*. Gaps in all the genome assemblies were identified using the ‘find_gaps’ by using the –fasta option and visualized in combination with the LOC126875579 klumps through the ‘klump_plot’ Klumpy subprogram.

## Software availability

The Klumpy Python package is available for download via ‘pip’ from the Python Package Index (PyPI). Source code for the package is available at https://bitbucket.org/Gio12/klumpy/src/master/. A script describing all steps in the analysis is available in Supplemental_Methods.pdf and at https://bitbucket.org/Gio12/klumpy/src/master/TestCases/TestCases.md.

## Acknowledgements

We thank Angel G. Rivera-Colón and Kira Long for their support and feedback during our project. This work was supported by the National Science Foundation (NSF OPP Grant 1645087 to JC).

