## Supplemental Material for "Klumpy: A Tool to Evaluate the Integrity of Long-Read Genome Assemblies and Illusive Sequence Motifs"

### Table of Contents

**Text S1** - Details of the Klumpy grouping algorithm.

**Table S1** - Lines of evidence used in Klumpy grouping algorithm.

**Figure S1** - Alignment plot of the canonical *afgp* locus in *C. gunnari*.

**Figure S2** - Alignment plot of the noncanonical *afgp* locus in *C. gunnari*.

**Table S2** - *Afgp* klumps identified in the *C. gunnari* reference genome.

**Figure S3** - Alignment plot of the canonical *afgp* locus in *C. esox*.

**Figure S4** - Alignment plot of the noncanonical *afgp* locus in *C. esox*.

**Table S3** - *Afgp* klumps identified in the *C. esox* reference genome.

**Figure S5** - Alignment plot of the *adcy5* region for the albino phenotype reference genome.

**Text S2** – Reconstruction of the *adcy5* gene neighborhood

**Figure S6** - Klump distribution for *lgi1*, *slc35g1*, *plce1*, *pcdh15*, *nmt2*, *kcnj3*, and *znf281* across the Illumina-based reference genome of the northern snakehead.

**Figure S7** - Klump distribution for *lgi1*, *slc35g1*, *plce1*, *pcdh15*, *nmt2*, *kcnj3*, and *znf281* across the PacBio-based reference genome of the northern snakehead.

**Figure S8** - Alignment of raw PacBio reads onto the unmapped locus at positions 26,726,550 – 26,739,233 on scaffold NW_026571047.1 using Mummer2 (Delcher et al. 2002).

**Figure S9** - Alignment plot of the great blue-spotted mudskipper Illumina paired-end data onto an unmapped region on NW_026571047.1.

**Figure S10** - Alignment plot of a contig generated using Flye (Kolmogorov et al. 2019) to reconstruct the unknown locus on scaffold NW_026571047.1.

**Figure S11** - Alignment plot illustrating the consistent tiling of raw reads up until the unmapped locus in the great blue-spotted mudskipper reference genome.

**Figure S12** - An alignment plot of the flagged region reported by Klumpy's genome scan on scaffold NC_066255.1 (positions 4.325 Mb – 4.4 Mb).

**Figure S13** - Klump plot of LOC126875579 klumps throughout the *Bombus huntii* reference genome.

**Figure S14** - Klump plot of LOC126875579 klumps throughout the *Bombus affinis* reference genome.

**Figure S15** - Klump plot of LOC126875579 klumps throughout the *Bombus hortorum* reference genome.

**Figure S16** - Klump plot of LOC126875579 klumps throughout the *Bombus hypnorum* reference genome.

**Figure S17** - Klump plot of LOC126875579 klumps throughout the *Bombus pratorum* reference genome.

**Figure S18** - Klump plot of LOC126875579 klumps throughout the *Bombus sylvestris* reference genome.

**Figure S19** - Klump plot of LOC126875579 klumps throughout the *Bombus terrestris* reference genome.

**Figure S20** - A flagged region in the *B. huntii* reference genome after implementing `scan_alignments` using default parameters.

### ***Text S1** – Details of the Klumpy grouping algorithm*

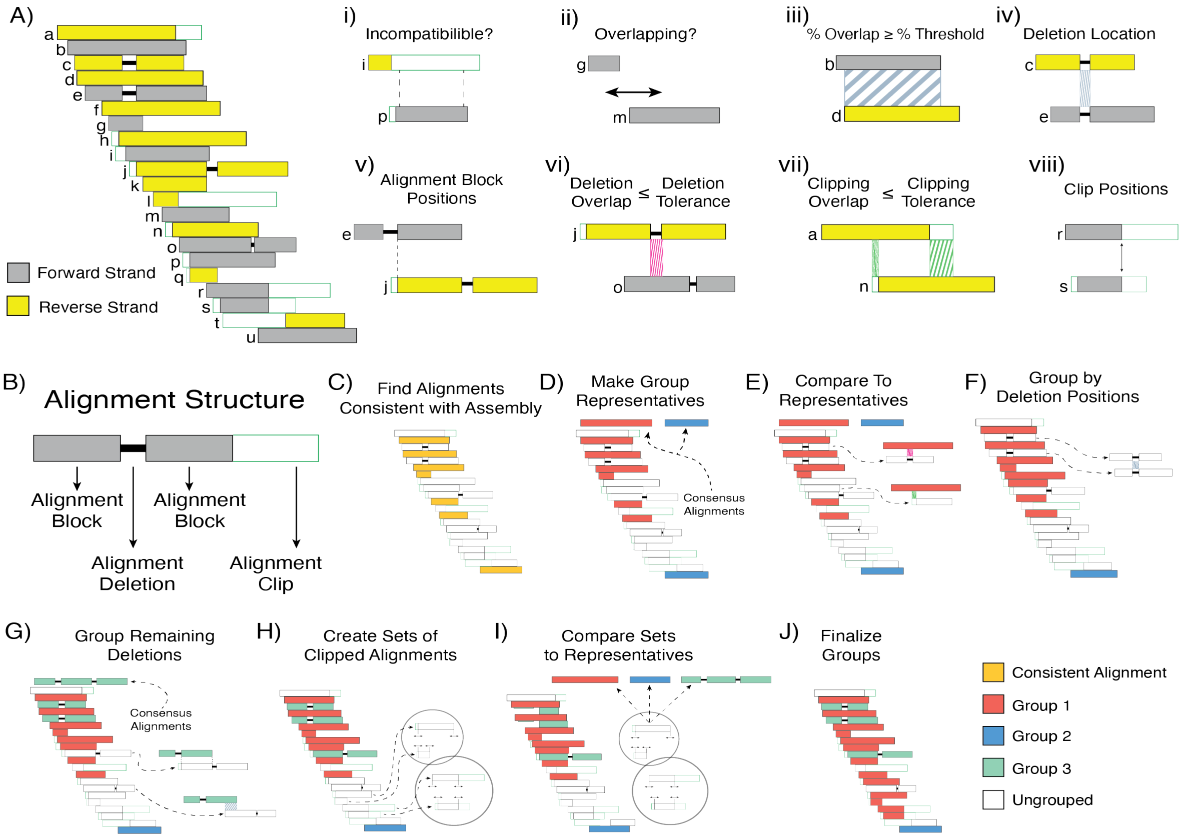


**Figure 6**: Alignment grouping algorithm. Given a set of alignments (A), each alignment's structure (B) is examined and used to collect pieces of evidence through a pairwise comparison (i-viii) to predict which alignments are derived from the same locus. An initial check for incompatibility is performed by checking whether one of the alignments falls into the clipped portion of the other alignment (i). If incompatibility is not overt, whether there is any overlap in aligned positions between the two alignments (ii) dictates the continuation of this process. Evidence is collected by checking the percentages of overlap between the two alignments (iii), observing whether the alignments contain deletions at the same positions (iv), inspecting the alignment block positions (v), examining the tolerance thresholds for alignment deletions and clips between the two alignments (vi, vii), and surveying the positions of any clipped sites (viii). To construct the groups, consistent alignments (C) are used to initialize the first group(s) and a representative alignment for each group (D). The algorithm then attempts to add the ungrouped alignments by comparing their structures to the consensus alignment(s) (E). If there remain inconsistent alignments containing deletions at the same positions, these alignments are then used to propose one or more new groups (F). A pairwise comparison across all alignments possessing at least one deletion is then performed to establish the proposed groups, which leads to the formation of one or more consensus alignments (G). Next, sets of unincorporated clipped alignments are assembled by reviewing the sites where the alignments are clipped (H). The sets of clipped sequences are then compared to the present groups for possible incorporation (I), at which point the grouping algorithm is terminated after removing groups not containing the minimum number of alignments required to form a group (J). A list of the lines of evidence along with their weights is shown in Table S1.

Here we define a *group* as a collection of sequences that share a similar alignment pattern. Given that many alignments may be inconsistent with the reference genome (e.g., soft or hard clipped, or extended via deletions), Klumpy takes into consideration several parameters that can be adjusted to determine if two alignments are able to be grouped together. In the initial grouping phase, consistent alignments (i.e., alignments that have no clipping or large deletions) are identified (Fig. 6K). As long-read sequencing is known to produce many indels (Carneiro et al. 2012; Jain et al. 2015, 2018), deletions that are shorter than the minimum deletion length are ignored. Simultaneously, alignments containing unignored deletions are recorded such that alignments with deletions in the same region in the window are associated with one another.

The initial groups are established by first identifying alignments consistent with the reference genome given its lack of clipped sites and inferred alignment deletions (Fig. 6C). A pairwise comparison amongst the consistent alignments is then performed to determine their level of compatibility (i.e., how likely are the two sequences to originate from the same locus). Compatibility scores are calculated for each pairwise comparison in order to obtain supporting evidence for grouping the alignments (Fig. 6i-6viii). To illustrate the compatibility scores, we will consider two alignments: *A* and *B*. If *A* is clipped at either end of the alignment and *B*'s aligned bases fall entirely in the clipped regions of *A*, the two sequences are considered incompatible (Fig. 6i). If *A* and *B* do not have any overlapping aligned bases, Klumpy considers it possible to group the two sequences together as there is no current evidence to suggest incompatibility nor compatibility (Fig. 6ii). Otherwise, the percentages *A* and *B* overlap each other in respect to their aligned positions, is computed (Fig. 6iii).

In cases where *A* and *B* both contain deletions, if the deletions occur at the same location, *A* and *B* are presumed to belong to the same group (Fig. 6iv). If the deletions do not agree but have alignment blocks (aligned bases separated by deletions) that begin or end at the same positions (e.g., if the second alignment block of *A* begins at the same position as the first alignment block of *B*), then the two sequences can also be grouped (Fig. 6v). The two sequences are considered incompatible in situations where the deletion from *A* overlaps more nucleotides in *B* than the set deletion tolerance (number of positions of the focal alignment overlapping a deletion in the other alignment) or vice versa (Fig. 6vi).

After examining any deletions contained in and between *A* and *B*, the algorithm then evaluates the clipping statuses of both alignments. If the number of clipped positions of *A* overlapping *B* is below the clipping tolerance (number of positions of the focal alignment overlapping a clipped region in the other alignment) and the case holds true for *B* (Fig. 6vii), the two alignments are then determined to belong to the same group in the case that the percentage overlap is above the minimum overlap percentage for both alignments. If not, the overlapping percentages is only considered as further evidence for grouping. If the grouping status between *A* and *B* has yet to be established (Fig. 6viii).

If the alignments start and are terminated at the same site in the reference genome, the alignments are considered to likely represent the same locus. If the number of deletions is the same between *A* and *B* and the total length of deleted positions is also identical, the two sequences are designated as part of the same group. In the case in which klumps are provided in the analysis when employing the `alignment_plot` subprogram, if there is no difference in the query sources found between *A* and *B*, an additional piece of evidence is obtained. Once these lines of evidence are collected and the two alignments are yet to be considered part of the same group, *A* and *B* are viewed as likely to belong to the same group if a value of 10 or more based on the supporting evidence is reached, reasonable if the value is at least 6, or unlikely otherwise (Table S1).

**Table S1.** Lines of evidence suggesting the grouping of two alignments. Attributes between the alignments are not equally weighted, as some realized criteria were seen to be more credible than others. Additionally, some lines of evidence are conditional on others (e.g., if two alignments begin at the same nucleotide, no further evidence is gained due to the alignments starting in close proximity to each other).

| Line of Evidence | Weight |
|---|---|
| 5’ end of one alignment overlaps the other alignment | +1 |
| 3’ end of one alignment overlaps the other alignment | +1 |
| Both alignments overlap one another no less than the minimum threshold | +1 |
| Both alignments begin at the same position | +2 |
| Both alignments begin within close proximity of each other | +1 |
| Both alignments end at the same position | +2 |
| Both alignments end within close proximity of each other | +1 |
| Both alignments have the same number of deletions | +1 |
| The total lengths of the deletions between the two alignments are equal | +1 |
| Alignments lacking deletions have tolerable clip overlaps with each other | +1 |
| The beginnings of the alignments have the same clipping status | +1 |
| The ends of the alignments have the same clipping status | +1 |
| Same klump sources are present in the alignments | +1 |

When performing a pairwise comparison amongst the consistent alignments, if one of the alignments overlaps the other alignment with a percentage equal to or greater than the minimum threshold (e.g., if 50% of one the alignments overlap with the other alignment), then the two alignments can be grouped together if the two alignments do not show incompatibility with the other alignments in the group under consideration. After initiating the groups with the consistent alignments, a consensus alignment is constructed for each group by merging the group’s alignment intervals to represent the group. (Fig. 6D). The ungrouped alignments are then compared to the consensus alignments, where sequence alignments that completely are overlapped by consensus alignment are incorporated in the group if any clipped or deletion lengths are below the tolerable thresholds (Fig. 6E). Returning to the ungrouped alignments that contain deletions, one or more groups are proposed based on the position of the deletions. Next, we iterate through the same alignments to ensure compatibility within the group and add additional overlapping alignments to form the groups ensuring the minimum number of alignments to form a group is held (Fig. 6F-G). The compatibility score check can be bypassed under the assumption that alignments should be grouped when deletions the overlap each other at the same genomic location and are tolerable with one another should be grouped.

Moving onto the alignments containing clips, the alignments are compared to one another to determine whether the alignments begin or end in close proximation of one another (Fig. 6H). Clipped alignments that are found to be anchored to similar positions are clustered into a set. This approach is founded on the intuition that alignments that are clipped at approximately the same position, are derived from a common locus. Once completed, sets where the number of alignments is below the minimum number of alignments to form a group are removed. Following, retained alignments are compared to the consensus alignments with the modification of allowing the clipped alignments to partially overlap with the consensus alignments. If compatible, the alignment and its corresponding set is added to the group, else a new group is formed (Fig. 6I).

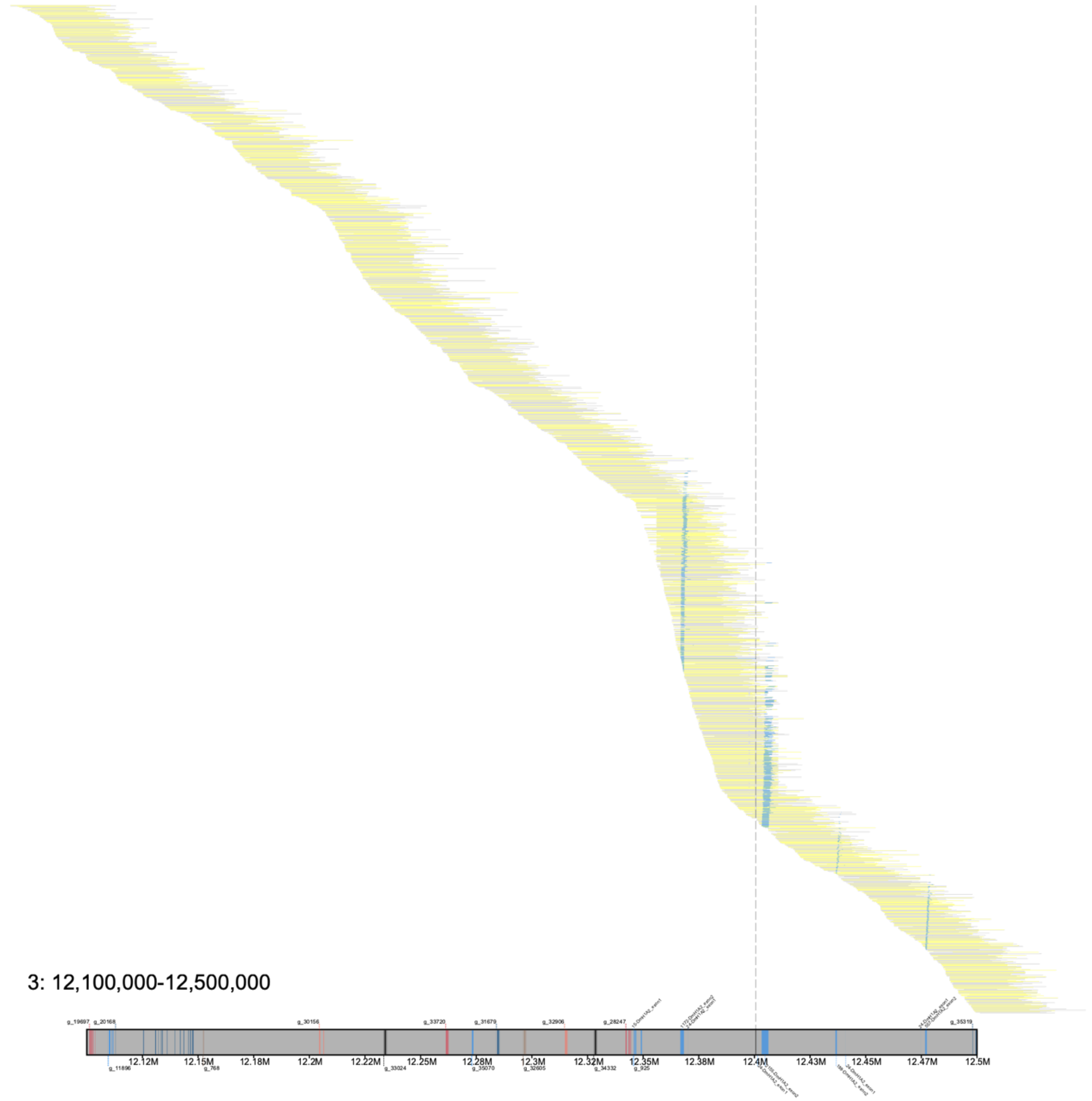


**Figure S1**: An alignment plot of the canonical *afgp* locus in *C. gunnari* on chromosome 3. Yellow blocks illustrate reads mapping onto the reverse orientation of the reference genome, while grey blocks represent alignments onto the forward orientation. See Fig. 2B for alignment structure depiction. Here, alignments were filtered to retain records with sequence lengths of at least 25 kilobases (Kbp), with a minimum of 75% of the base pairs being mapped. *Afgp* klumps are drawn on both the reference genome and the aligned sequences (light blue boxes), with the reference klumps labeled with *N*-DmH1A2_exon*M*, where *N* is the number of k-mers in the klump, and *M* is indicating which exon (exon 1 or exon 2) the klump is derived from. A gap in the assembly is illustrated with a dashed vertical grey bar. Notably, the alignments are able to span across the gap, providing evidence for a correct assemblage of this locus. The remaining annotations on the reference genome symbolize the surrounding genes based on the gene annotations of the assembly.

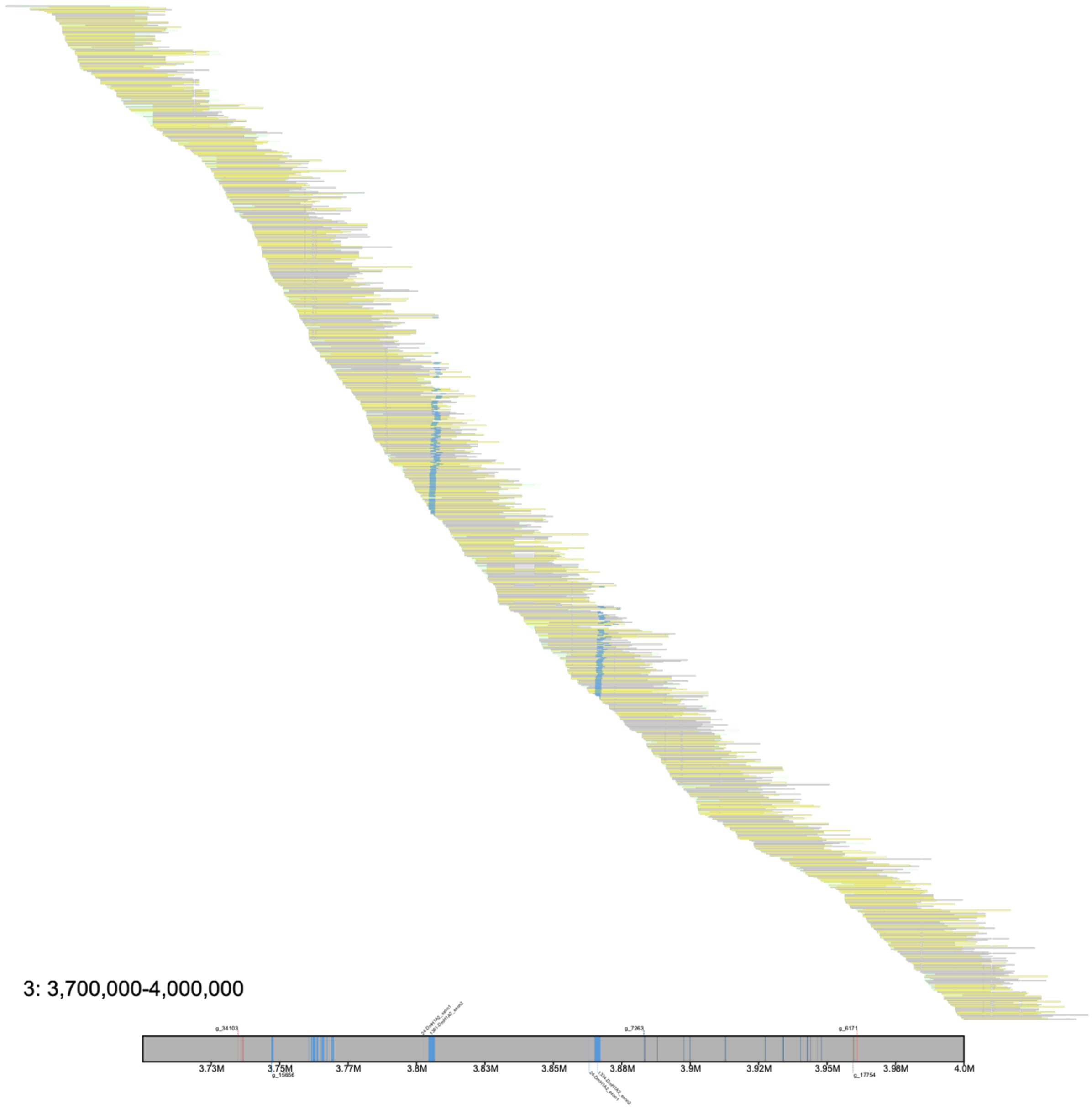


**Figure S2**: An alignment plot of the noncanonical *afgp* region in *C. gunnari* between 3.7 Mb and 4 Mb on chromosome 3. Visuals follow the same scheme as those describe in Fig. S1. Aligned sequences illustrated are 25 Kbp or greater in length and have no more than a quarter of their base pairs unmapped. A notable feature is the lack of gaps in the locus and the consistent tiling of the alignments, which confirm the assemblage of this region.

**Table S2**: *Afgp* klumps identified in the *C. gunnari* reference genome. Klumps were generated using a maximum distance of 1 Kbp between k-mers and a minimum klump size of 10 k-mers. DmH1A2_exon1 and DmH1A2_exon2 were the sequences used for the first and second exon for the *afgp* gene model obtained from Nicodemus-Johnson et al. (2011). K-merizing the first exon with a k-mer size of 17 results in 24 k-mers, and thus we consider a complete copy of the *afgp* exon 1 sequence being partially characterized with a klump of 24 k-mers.

| Chromosome | Start Position | End Position | Number of k-mers | Query Source | Orientation |
|---|---|---|---|---|---|
| 3 | 3802399 | 3802438 | 24 | DmH1A2_exon1 | F |
| 3 | 3863088 | 3863127 | 24 | DmH1A2_exon1 | F |
| 3 | 12345701 | 12345731 | 15 | DmH1A2_exon1 | F |
| 3 | 12401363 | 12401402 | 24 | DmH1A2_exon1 | F |
| 3 | 12474642 | 12474681 | 24 | DmH1A2_exon1 | F |
| 3 | 3804368 | 3806620 | 1361 | DmH1A2_exon2 | F |
| 3 | 3865048 | 3867220 | 1334 | DmH1A2_exon2 | F |
| 3 | 12403237 | 12406314 | 2155 | DmH1A2_exon2 | F |
| 3 | 12476694 | 12477486 | 557 | DmH1A2_exon2 | F |
| 3 | 12436393 | 12437075 | 188 | DmH1A2_exon2 | R |
| 3 | 12366688 | 12368315 | 1172 | DmH1A2_exon2 | R |
| 3 | 12440970 | 12441009 | 24 | DmH1A2_exon1 | R |
| 3 | 12370174 | 12370213 | 24 | DmH1A2_exon1 | R |

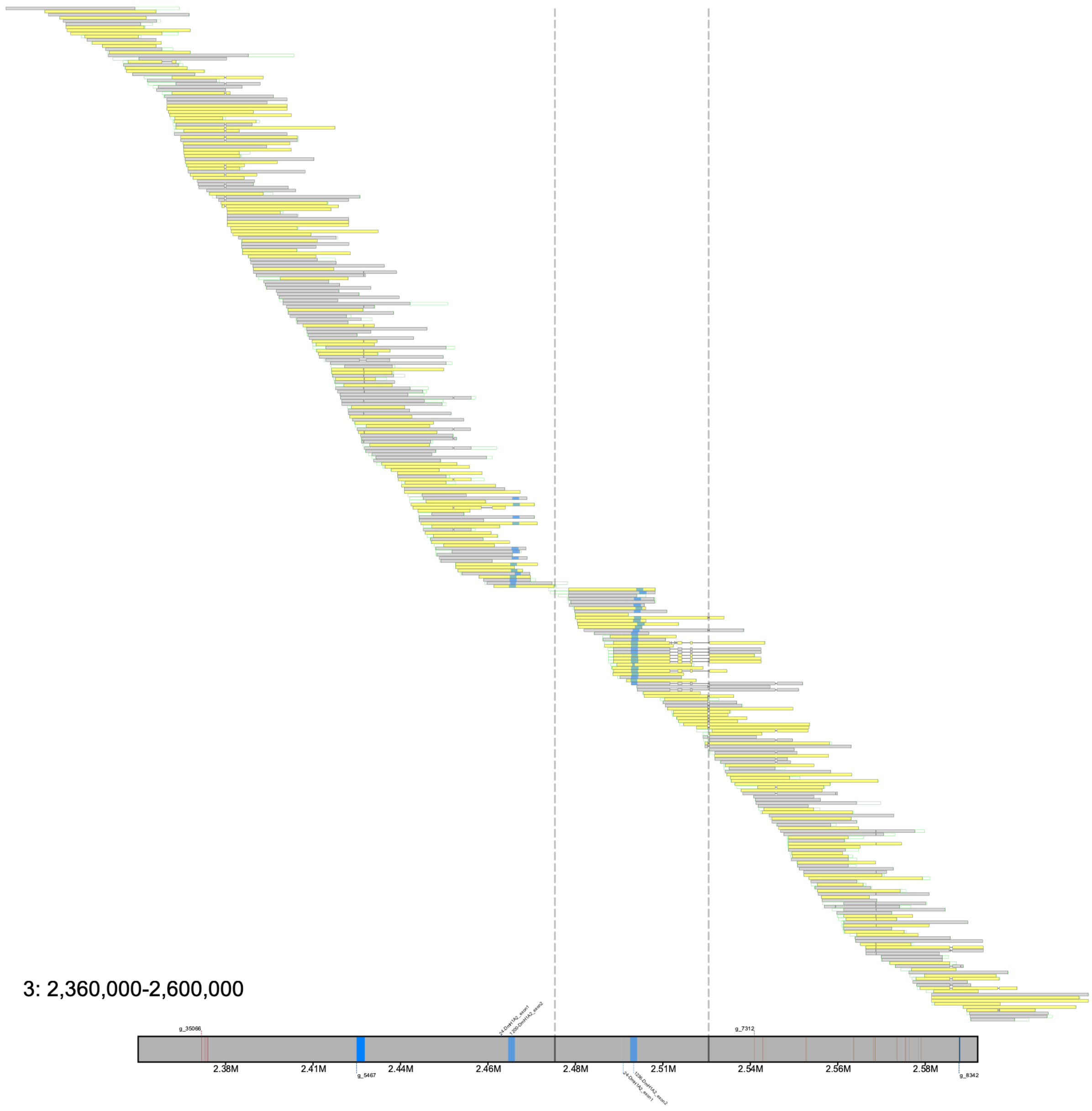


**Figure S3**: Alignment plot of the noncanonical *afgp* locus on chromosome 3 for *C. esox*. See the caption of Fig. S1 for a description regarding the annotations and properties of the image. Aligned reads have at least 75% its bases mapped and are equal or greater than 15 Kbp in length. Light blue boxes illustrate the *afgp* klumps found in the reference genome and aligned reads. Two grey dashed lines illustrate two gaps in the assembly; however, reads mapping between the middle and rightmost contig are able to span the gap, indicating a proper merging of the two contigs. Based on gene neighborhoods between *C. gunnari* and *C. esox*, the leftmost contig is properly placed in order to maintain conserved synteny between the two species (Rivera-Colón et al. 2023).

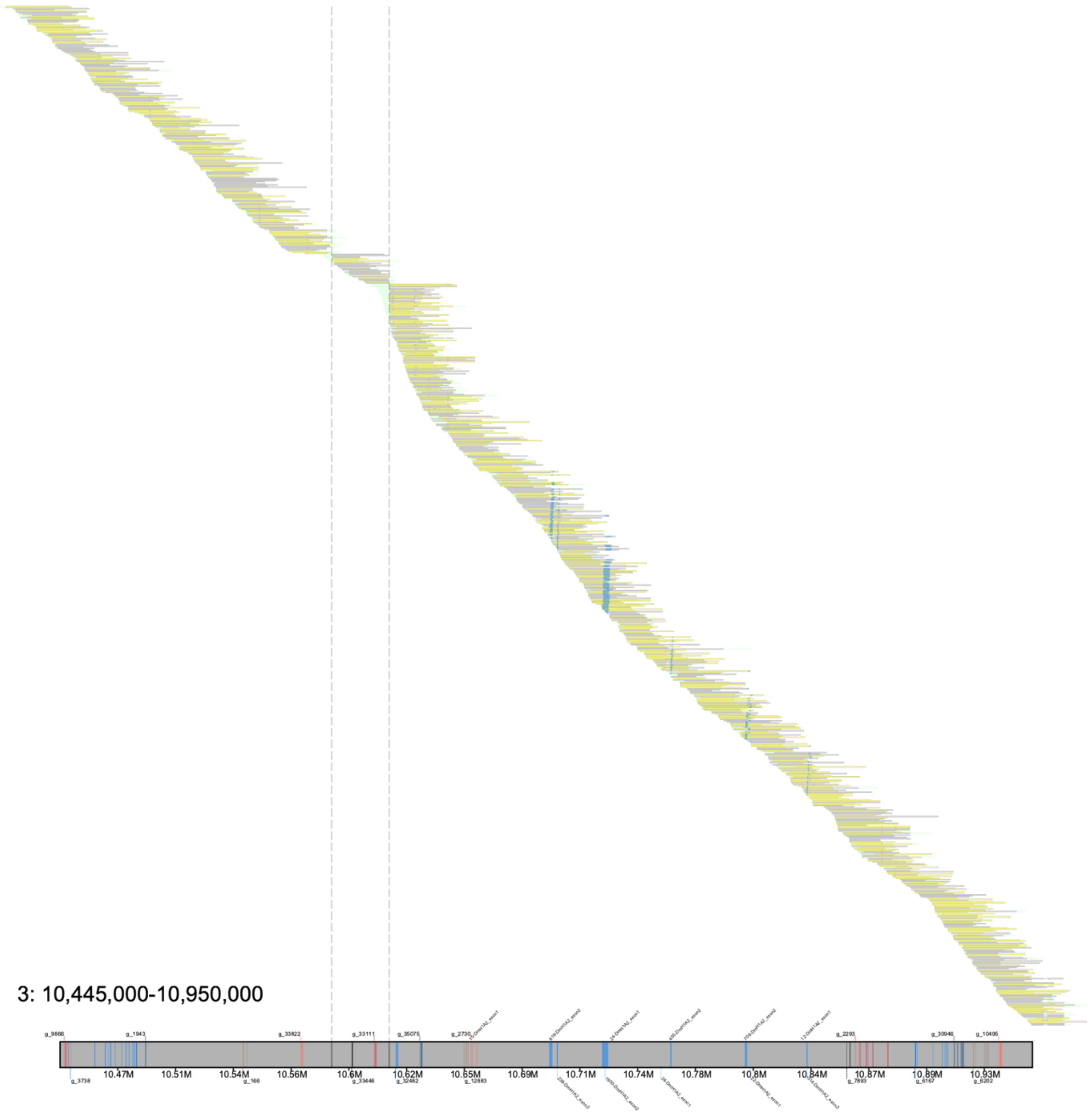


**Figure S4**: Alignment plot of the canonical *afgp* locus on chromosome 3 for *C. esox*. Descriptions for the features in the image can be found in Fig. S1. In the illustration, alignments with reads shorter than 15 Kbp and with more than 25% of the bases unmapped are not included. Between 10.445 Mb and 10.95 Mb, three contigs are united by gaps between 10.56 Mb and 10.62 Mb, and despite of lack of alignments spanning across the gaps, a synteny-based approach was implemented to support this region (Rivera-Colón et al. 2023). Consistent tiling of the alignments for all three contigs does not present any evidence for any irregularities in the assembly.

**Table S2**: *Afgp* klumps identified in the *C. esox* reference genome. Klumps were generated following the same procedure described in the caption of Table S1.

| Chromosome | Start Position | End Position | Number of k-mers | Query Source | Orientation |
|---|---|---|---|---|---|
| 3 | 2463765 | 2463804 | 24 | DmH1A2_exon1 | F |
| 3 | 2498651 | 2498690 | 24 | DmH1A2_exon1 | F |
| 3 | 10658240 | 10658270 | 15 | DmH1A2_exon1 | F |
| 3 | 10757057 | 10757096 | 24 | DmH1A2_exon1 | F |
| 3 | 10830707 | 10830734 | 12 | DmH1A2_exon1 | F |
| 3 | 2465773 | 2467719 | 1200 | DmH1A2_exon2 | F |
| 3 | 2500652 | 2502656 | 1236 | DmH1A2_exon2 | F |
| 3 | 10702947 | 10703573 | 238 | DmH1A2_exon2 | F |
| 3 | 10761826 | 10762599 | 455 | DmH1A2_exon2 | F |
| 3 | 10832651 | 10833212 | 314 | DmH1A2_exon2 | F |
| 3 | 10800628 | 10801828 | 709 | DmH1A2_exon2 | R |
| 3 | 10726465 | 10729658 | 1800 | DmH1A2_exon2 | R |
| 3 | 10699091 | 10700701 | 619 | DmH1A2_exon2 | R |
| 3 | 10803629 | 10803666 | 22 | DmH1A2_exon1 | R |
| 3 | 10731475 | 10731514 | 24 | DmH1A2_exon1 | R |

*Searching for the adenylate cyclase 5 (adcy5) gene in the northern snakehead*

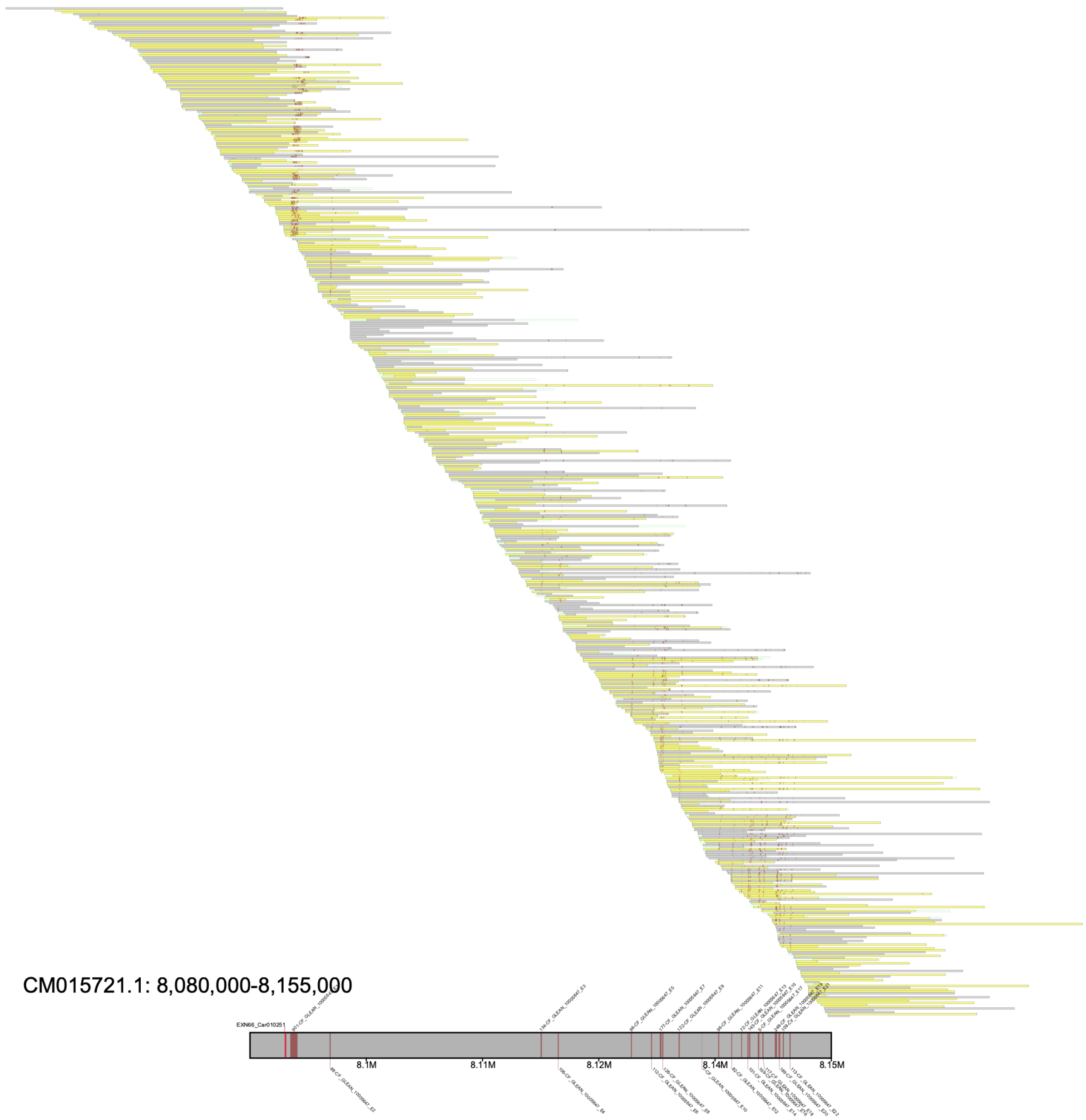


**Figure S5**: Alignment plot of the *adcy5* region for the albino phenotype reference genome. See the caption of Fig. S1 for a description regarding the annotations and properties of the image. Alignments in which with more than half the sequence being unmapped or have a sequence length less than 2 Kbp were excluded from the plot. Maroon blocks on the reference genome and raw reads illustrate *adcy5* klumps. The consistent tiling of the alignments across this gapless locus and the *adcy5* klumps detected fail to suggest a misassembled locus as a potential source of an incomplete annotation of *adcy5* in the reference genome. Of note, upstream of the first *adcy5* klump is the *adcy5* annotation from Zhou et al. (2022).

***Text S2*** *– Reconstruction of adcy5 gene neighborhood*

In a study conducted by Zhou et al. (2022), the authors reported the absence of the adenylate cyclase 5 (*adcy5*) gene in the genome assembly of the albino northern snakehead (*Channa argus*). The researchers outline the gene neighborhood for the *adcy5* gene for both the albino and non-albino northern snakehead reference genomes. More specifically, they present genes *lgi1*, *slc35g1*, *plce1*, and *pcdh15* downstream of where the *adcy5* gene would be located assuming collinearity between the two assemblies, and genes *nmt2*, *kcnj3*, and *znf281* upstream from the locus. After locating the *adcy5* gene (see main text), we sought to annotate the reported gene neighborhood.

To obtain gene models for the genes of interest (*lgi1*, *slc35g1*, *plce1*, *pcdh15*, *nmt2*, *kcnj3*, and *znf281*), we retrieved the exonic sequences from the climbing perch (*Anabas testudineus*) reference assembly (Rhie et al. 2021) through Ensembl (Cunningham et al. 2022). The sequences were then used as queries in Klumpy's subprogram `find_klumps` to generate klumps by setting the minimum klump size to 4 k-mers, a max distance of 10 bp and retaining sequences containing klumps from at least 5 different queries. Using the same approach, *adcy5* klumps for the Illumina-based (i.e., the non-albino phenotype) assembly were generated using the assembly's *adcy5* exons. A description on the detection of *adcy5* klumps for the PacBio (i.e., the albino phenotype) assembly can be found in the main text. The klump distribution in the reference genomes for the non-albino phenotype and albino phenotype can be viewed in Figures S6 and S7, respectively.

Only three of the genes (*lgi1*, *slc35g1*, *plce1*) occupy the same gene neighborhood for both assemblies (Figures S6 and S7) as reported by Zhou et al. (2022). Moreover, this study is in concert with Sun et al. (2023), as there is no evident association between the albino phenotype and the *adcy5* gene. By implementing an automated pipeline, Klumpy demonstrates its utility to not only locate a set of query sequences but provide the user with a graphical result for a straightforward and quick comprehension of the results.

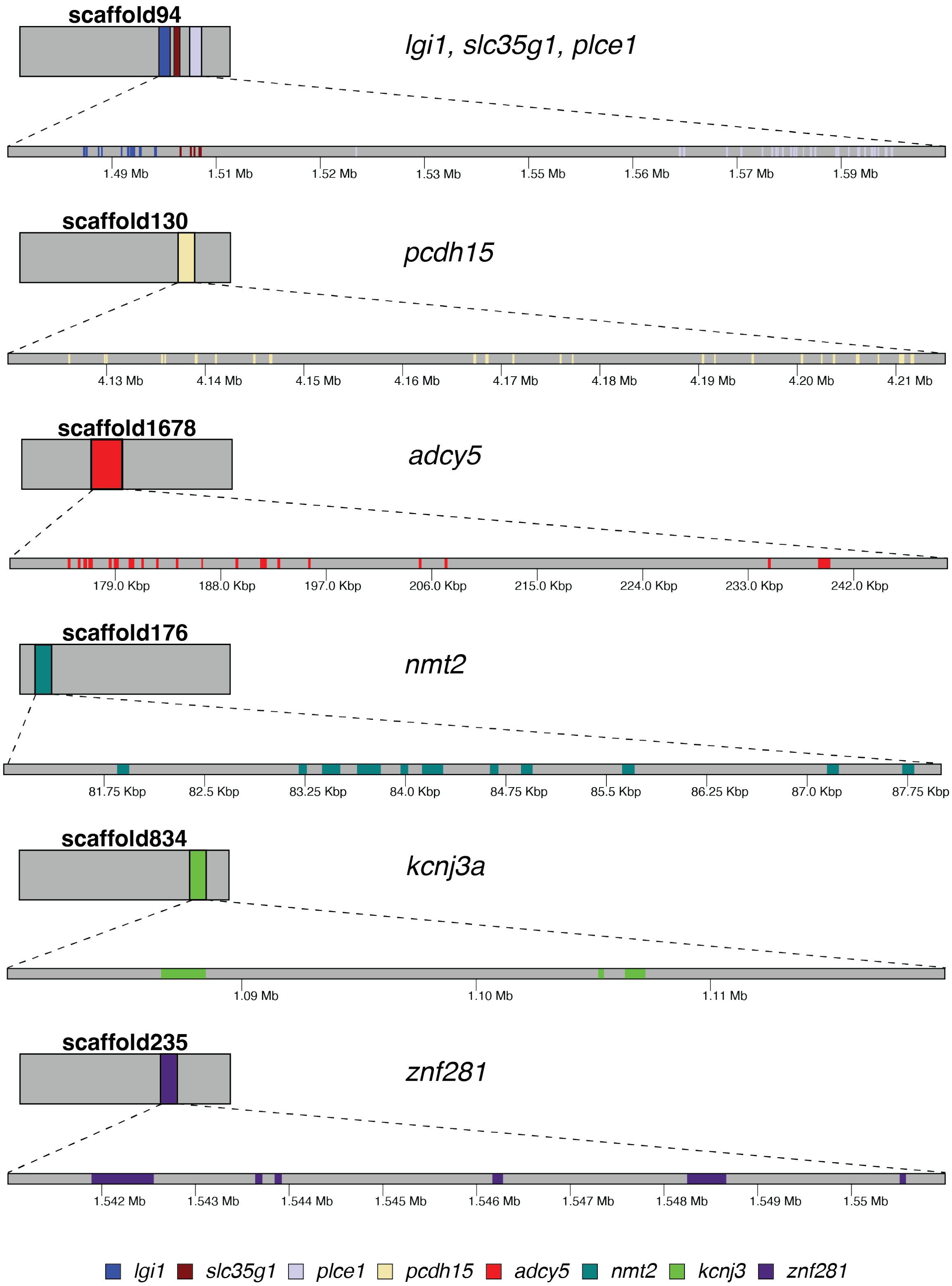


**Figure S6**: Klump distribution for *lgi1*, *slc35g1*, *plce1*, *pcdh15*, *nmt2*, *kcnj3*, and *znf281* across the Illumina-based reference genome of the northern snakehead. Genes are located across six separate scaffolds. Shorter rectangles represent entire contigs, with the longer rectangles portraying a zoomed in view of the identified klumps. Klump distribution figures were produced using the `klump_plot` subprogram and modified for clarity.

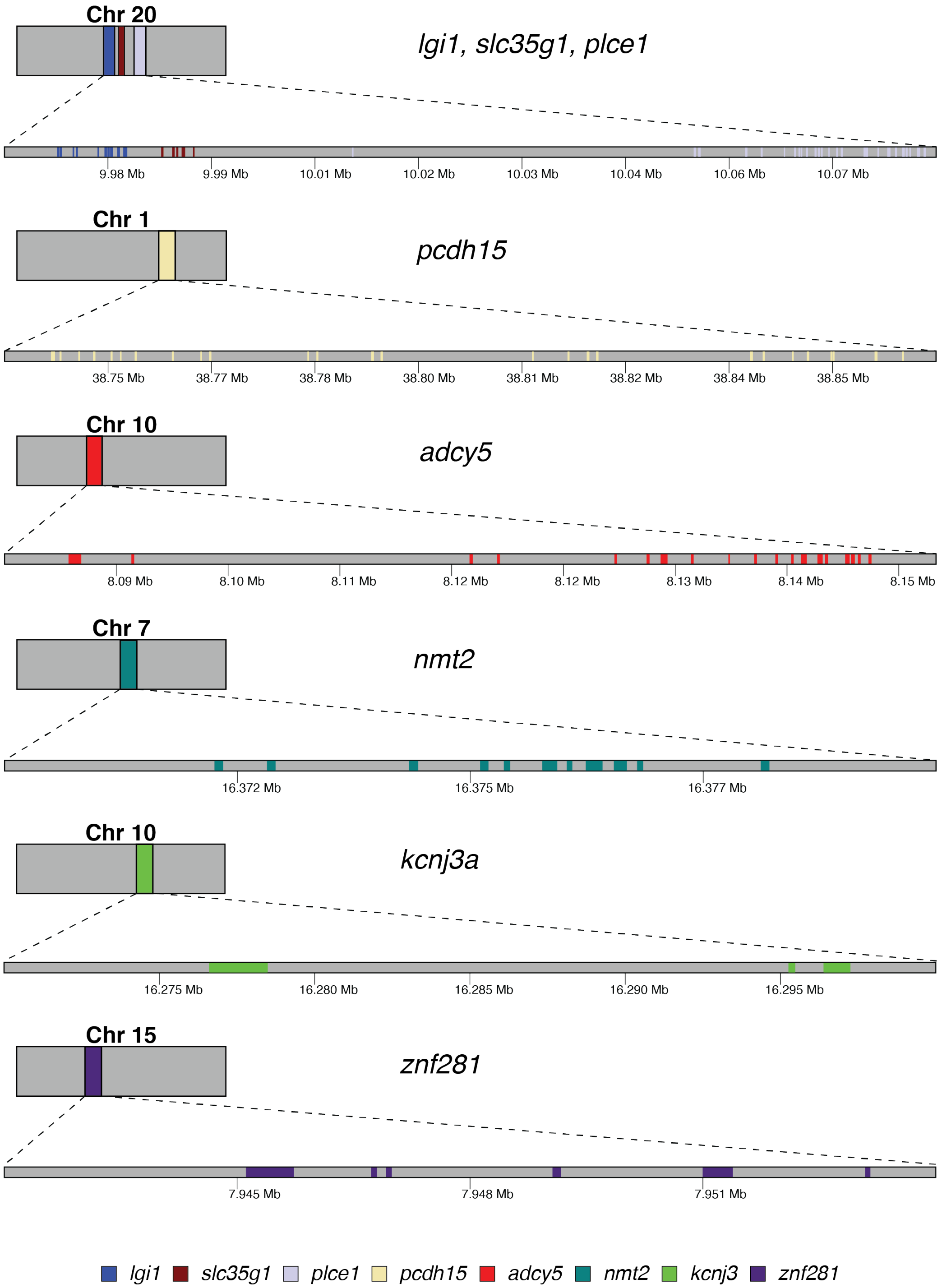


**Figure S7**: Klump distribution for *lgi1*, *slc35g1*, *plce1*, *pcdh15*, *nmt2*, *kcnj3*, and *znf281* across the PacBio-based reference genome of the northern snakehead. Genes are located across five chromonomes. Shorter rectangles represent entire scaffolds, with the longer rectangles displaying the located klumps in a closer illustration. Klump distribution figures were produced using the `klump_plot` subprogram and modified for clarity.

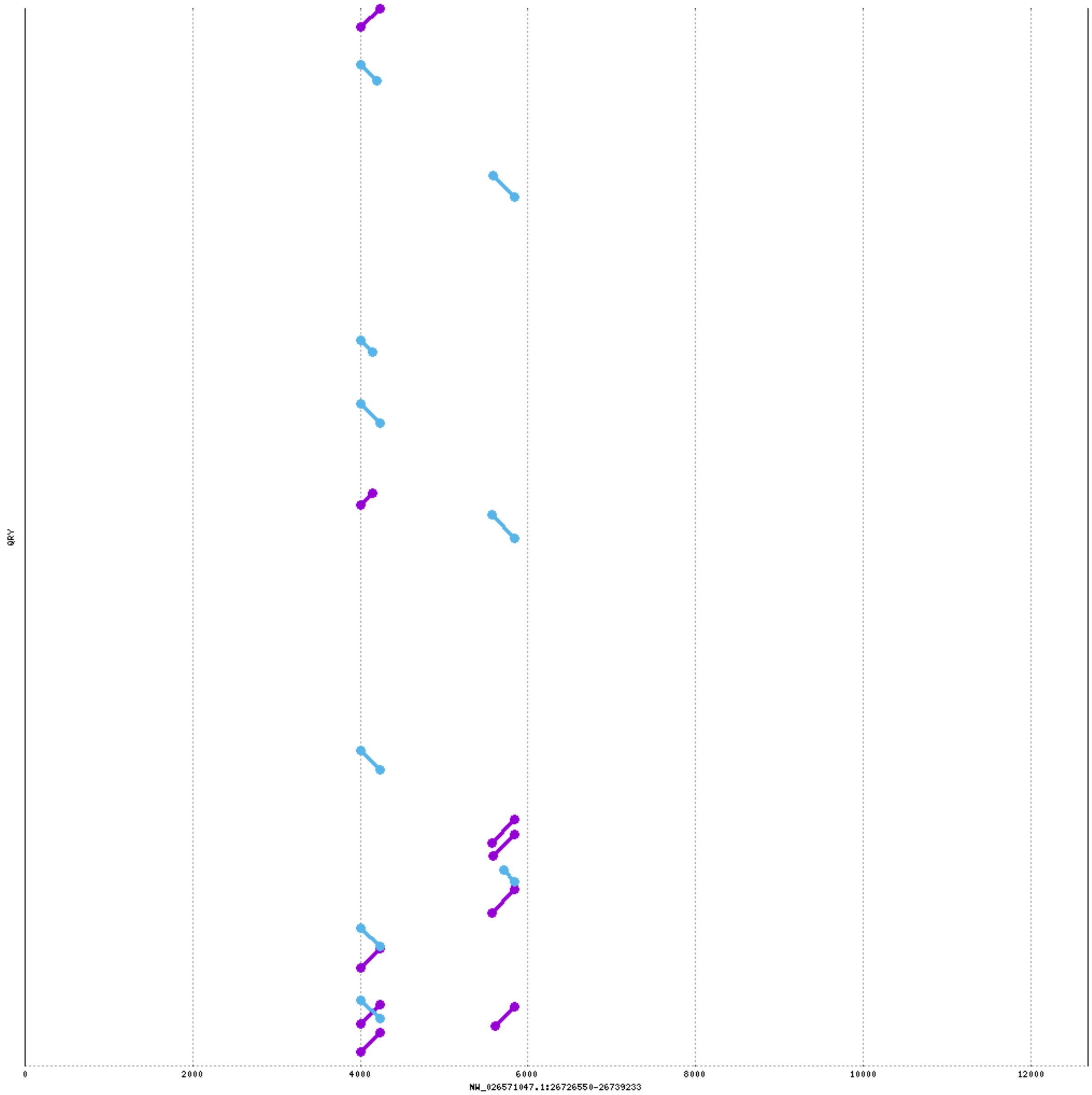


**Figure S8**: Alignment of raw PacBio reads onto the unmapped locus at positions 26,726,550 – 26,739,233 on scaffold NW_026571047.1 using Mummer2 (Delcher et al. 2002). Most of the locus remains uncovered, with a few matching k-mers (purple and blue connections) from the raw reads present at two sites within the region.

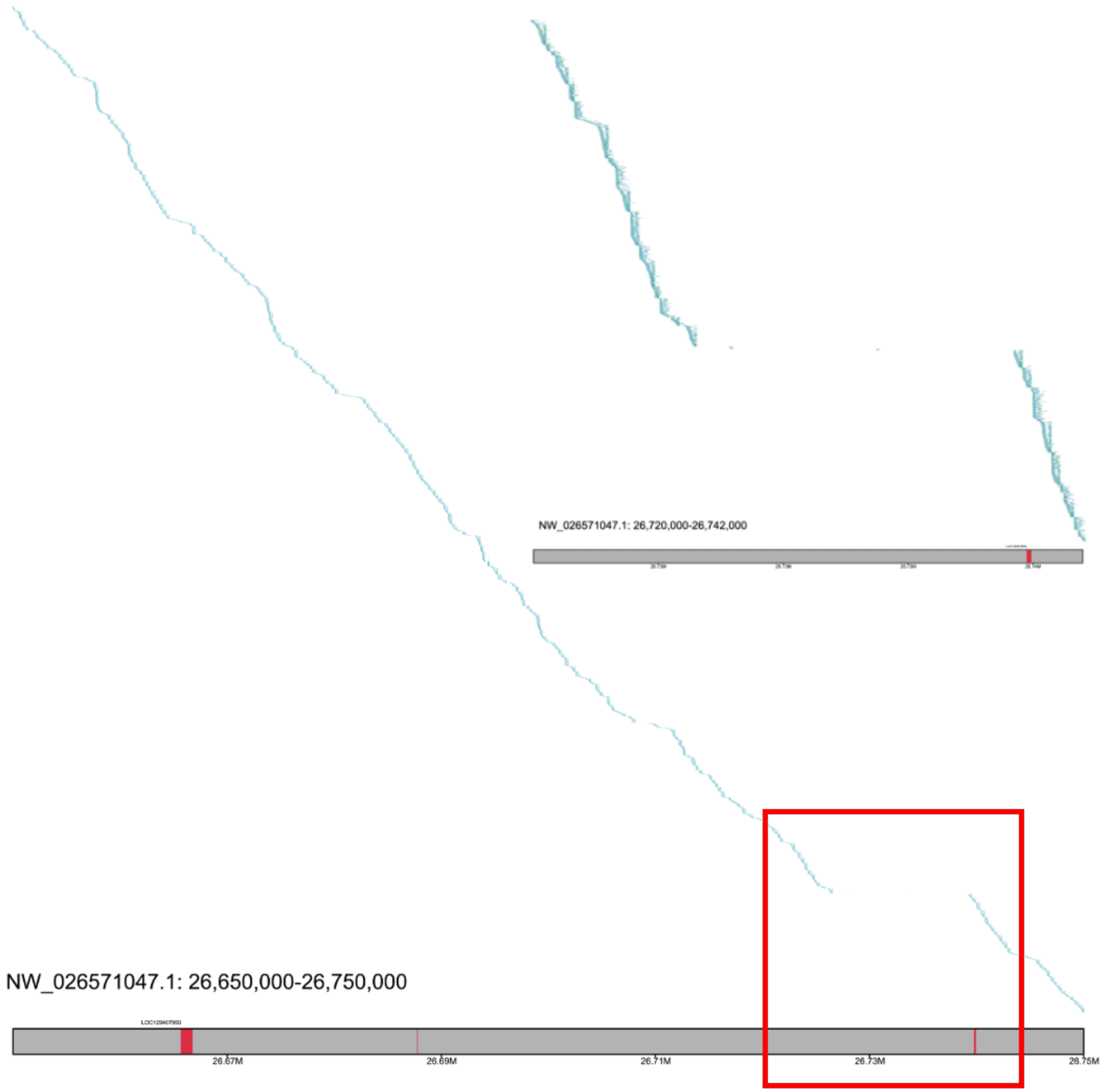


**Figure S9**: Alignment plot of the great blue-spotted mudskipper Illumina paired-end data onto an unmapped region on NW_026571047.1. Reads with their pair are mapped onto the same horizontal axis, with the first of the pair in a shade of green-cyan, and the second in the pair colored a cyan-blue. The annotation LOC129407950 (identified as *insyn2a*) is shown as red boxes within the reference genome. Alignments below a length of 100 bp and with less than half the read aligned, are excluded. The red unfilled box highlights a zoomed in portion of the alignment shown in the upper right of the figure. Similarly to the PacBio data (Fig. S11), the cryptic locus is largely unmapped. Alignments at two sites within the locus are comparable to the alignment of PacBio reads using Mummer2 (Delcher et al. 2002; Fig. S8).

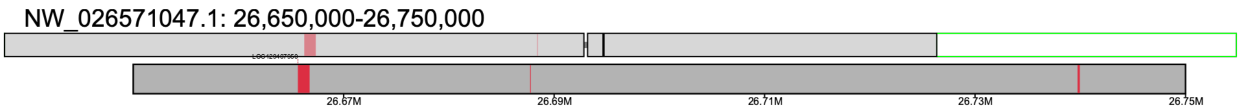


**Figure S10**: Alignment plot of a contig generated using Flye (Kolmogorov et al. 2019) to reconstruct the unknown locus on scaffold NW_026571047.1. A description for the interpretation of an alignment plot can be viewed in the caption of Fig. S1. Reads used to assemble this region were taken from alignments mapping to positions 26.65 Mb – 26.75 Mb at this scaffold. Both the reference assembly and contig contain *insyn2a* klumps (here, called LOC110174603); however, the contig is soft clipped (clear green rectangle) at the locus containing the ambiguous region.

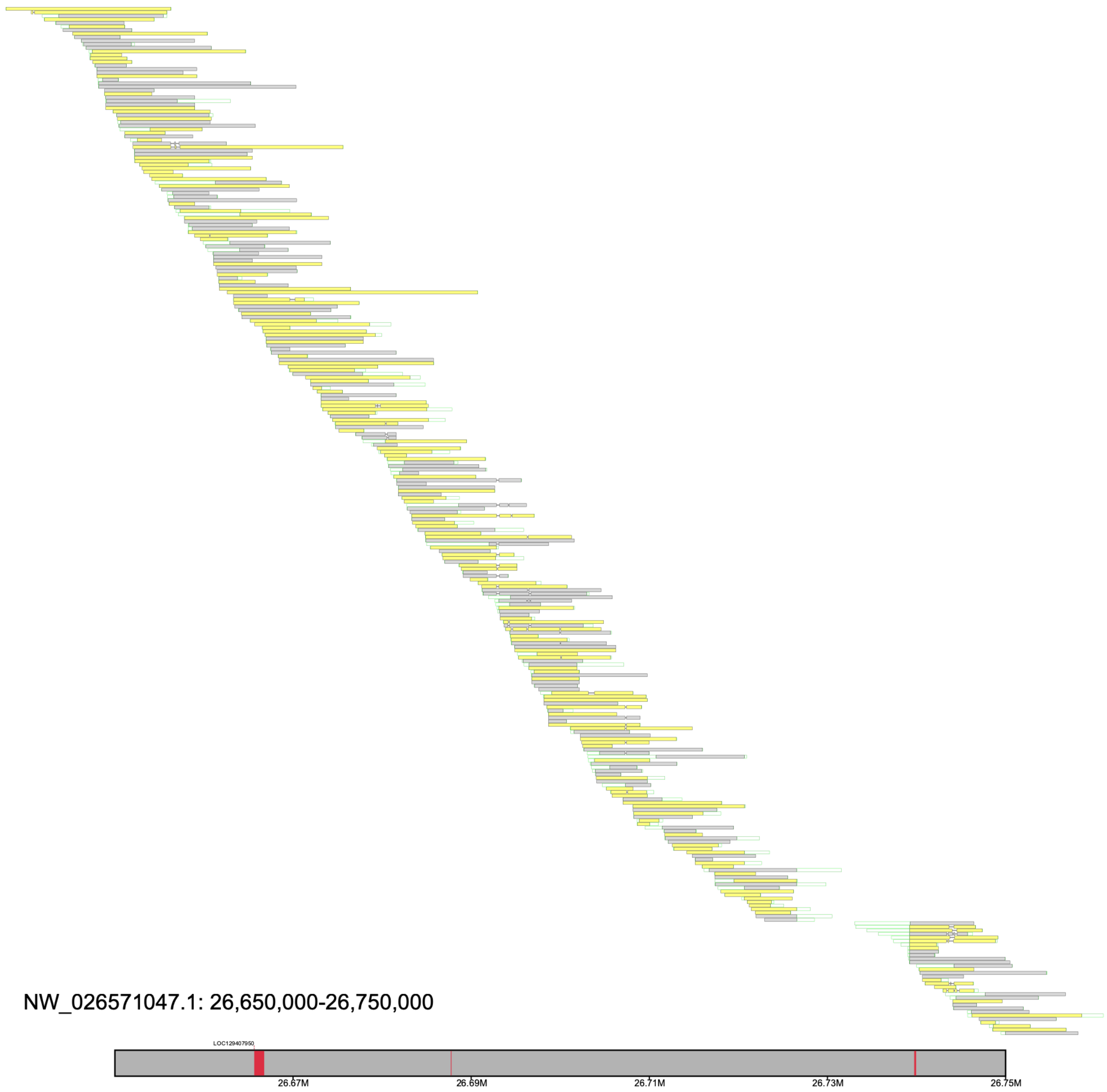


**Figure S11**: Alignment plot illustrating the consistent tiling of raw reads up until the unmapped locus in the great blue-spotted mudskipper reference genome. Alignments were filtered to retain alignments with at least half the sequence mapped. Features of the plot (e.g., yellow rectangles) are described in the Fig. S1 caption. Here, the red rectangles on the reference genome are representing the exons of LOC110174603, which was identified as the *insyn2a* gene. Alignments flanking the unmapped region are soft clipped (clear green rectangles) at approximately the same positions.

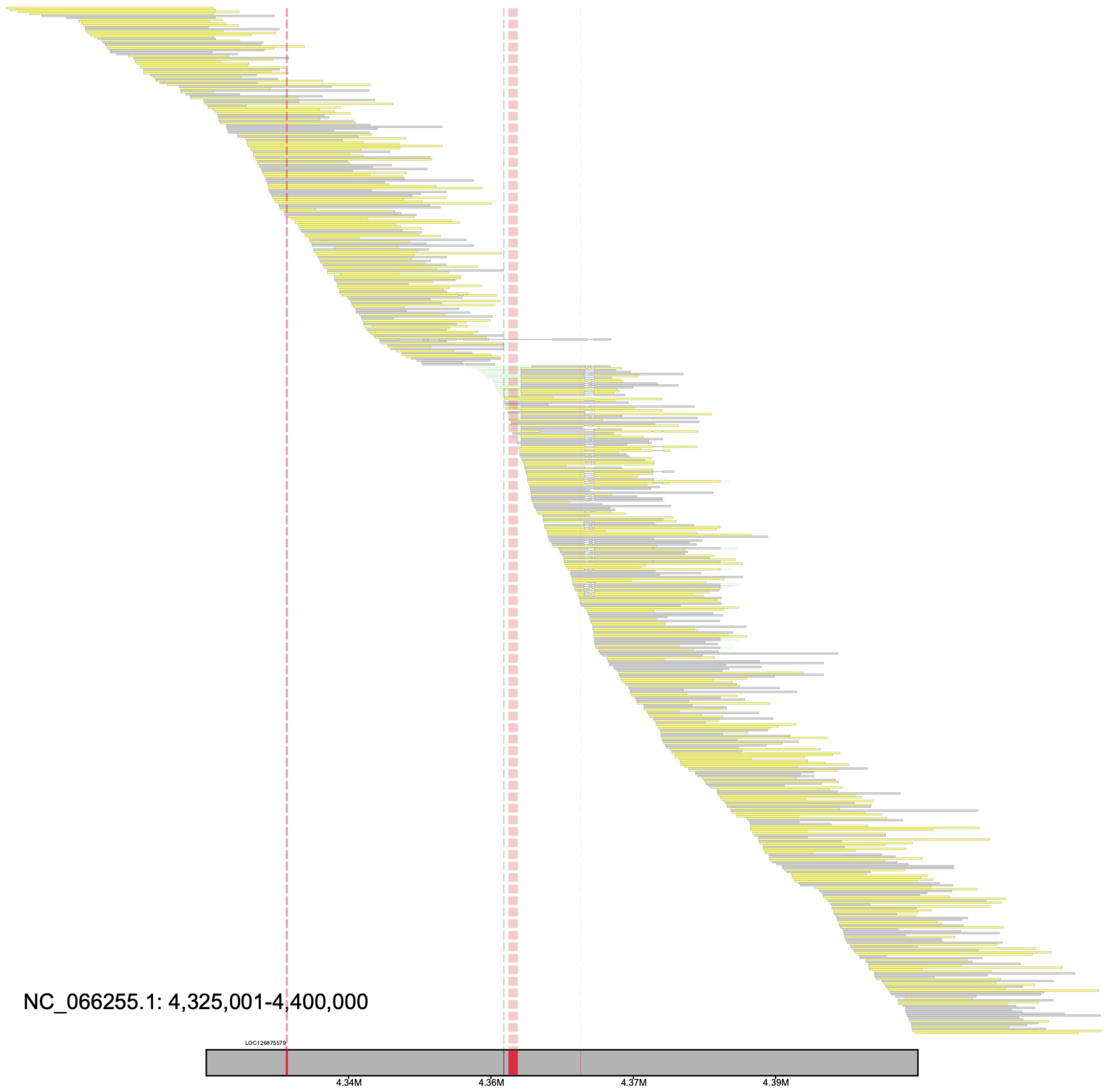


**Figure S12**: An alignment plot of the flagged region reported by Klumpy's genome scan on scaffold NC_066255.1 (positions 4.325 Mb – 4.4 Mb). Aligned records are grey if aligned to the forward orientation; other, they are yellow to indicate an alignment to the reverse orientation. Green clear boxes flanking the alignments are representative of soft clips. Within the window, the three exons for LOC126875579 are present in the assembly (red blocks in reference genome), with red dash lines highlighting their locations. Dashed grey line ~4.36 Mb illustrates a gap in the assembly, where two contigs were merged during the scaffolding process. Most alignments flanking the gap are soft clipped and unable to span the gap. Noticeably, the three exons for LOC126875579 are found across the two contigs. Moreover, many alignments are clipped at the positions corresponding to the second exon of LOC126875579.

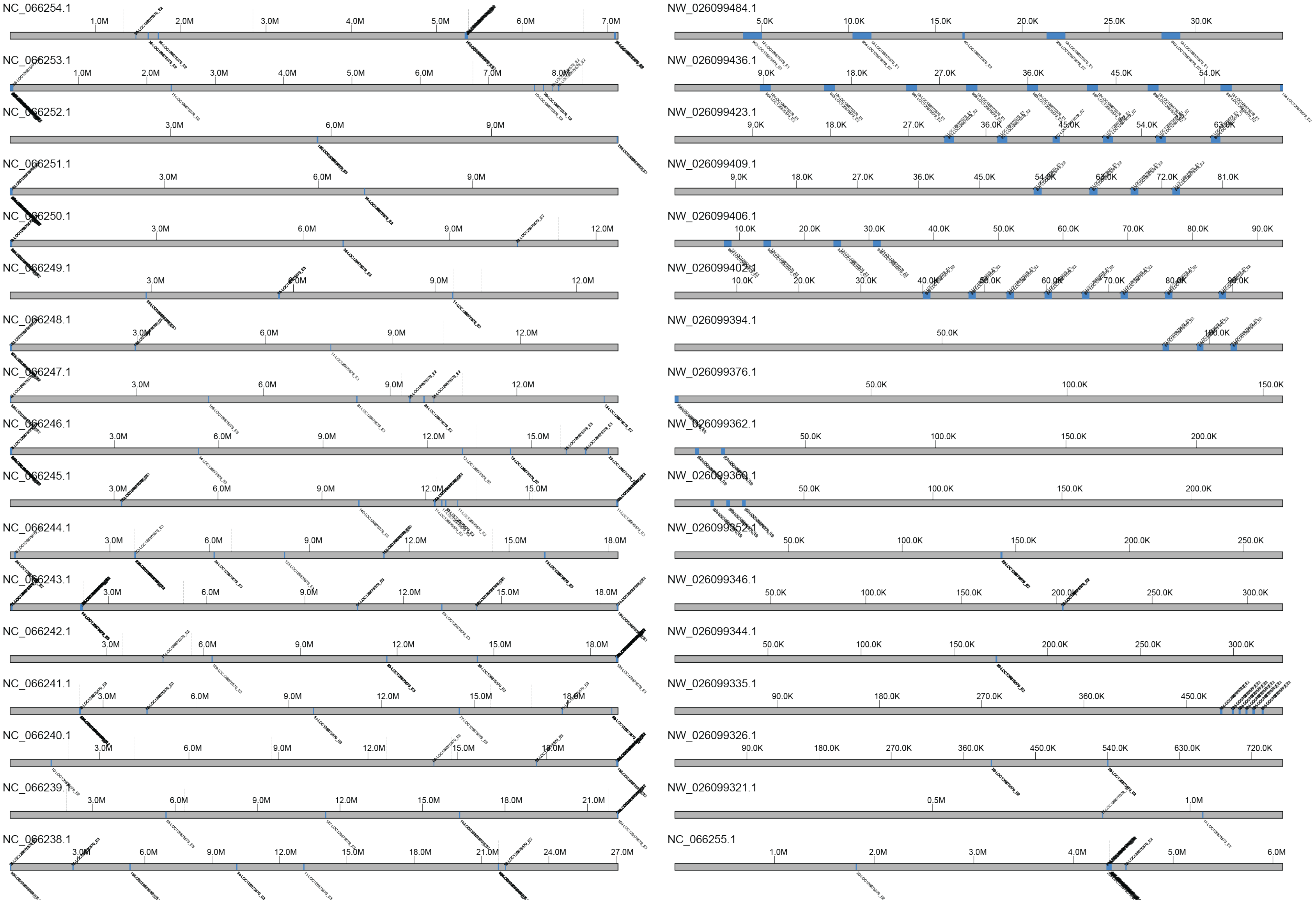


**Figure S13**: Klump plot of LOC126875579 klumps throughout the *Bombus huntii* reference genome. Blue boxes illustrate LOC126875579 klumps, and dashed lines above each sequence represent gaps in the sequence.

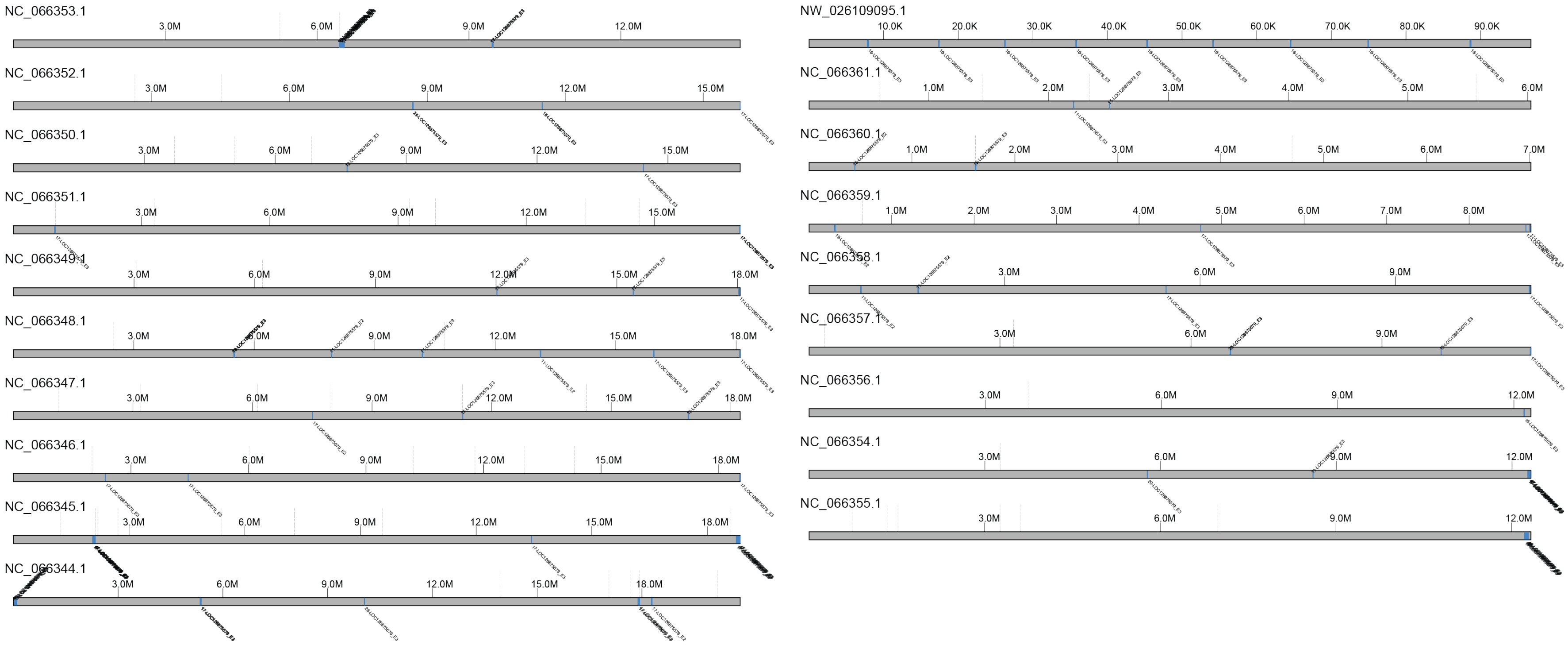


**Figure S14**: Klump plot of LOC126875579 klumps throughout the *Bombus affinis* reference genome. Blue boxes illustrate LOC126875579 klumps, and dashed lines above each sequence represent gaps in the sequence.

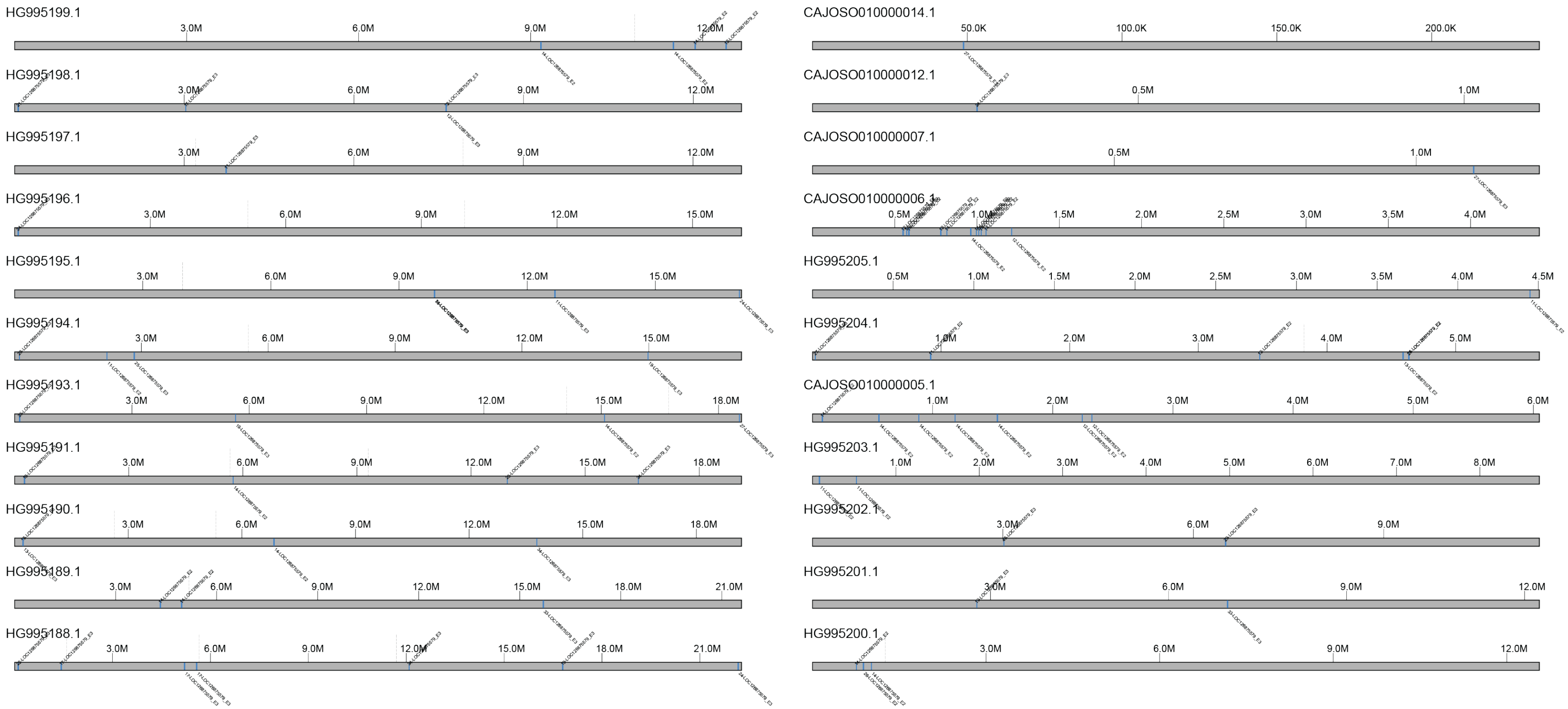


**Figure S15**: Klump plot of LOC126875579 klumps throughout the *Bombus hortorum* reference genome. Blue boxes illustrate LOC126875579 klumps, and dashed lines above each sequence represent gaps in the sequence.

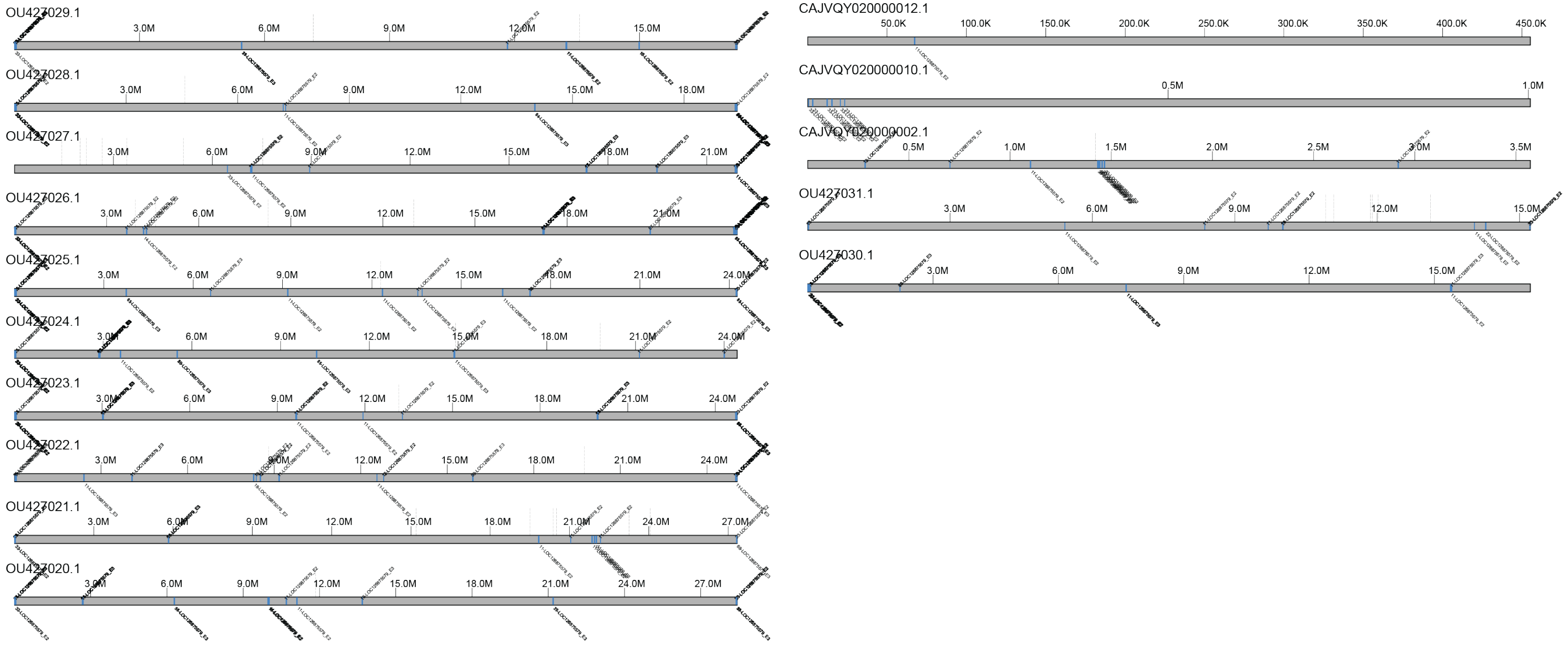


**Figure S16**: Klump plot of LOC126875579 klumps throughout the *Bombus hypnorum* reference genome. Blue boxes illustrate LOC126875579 klumps, and dashed lines above each sequence represent gaps in the sequence.

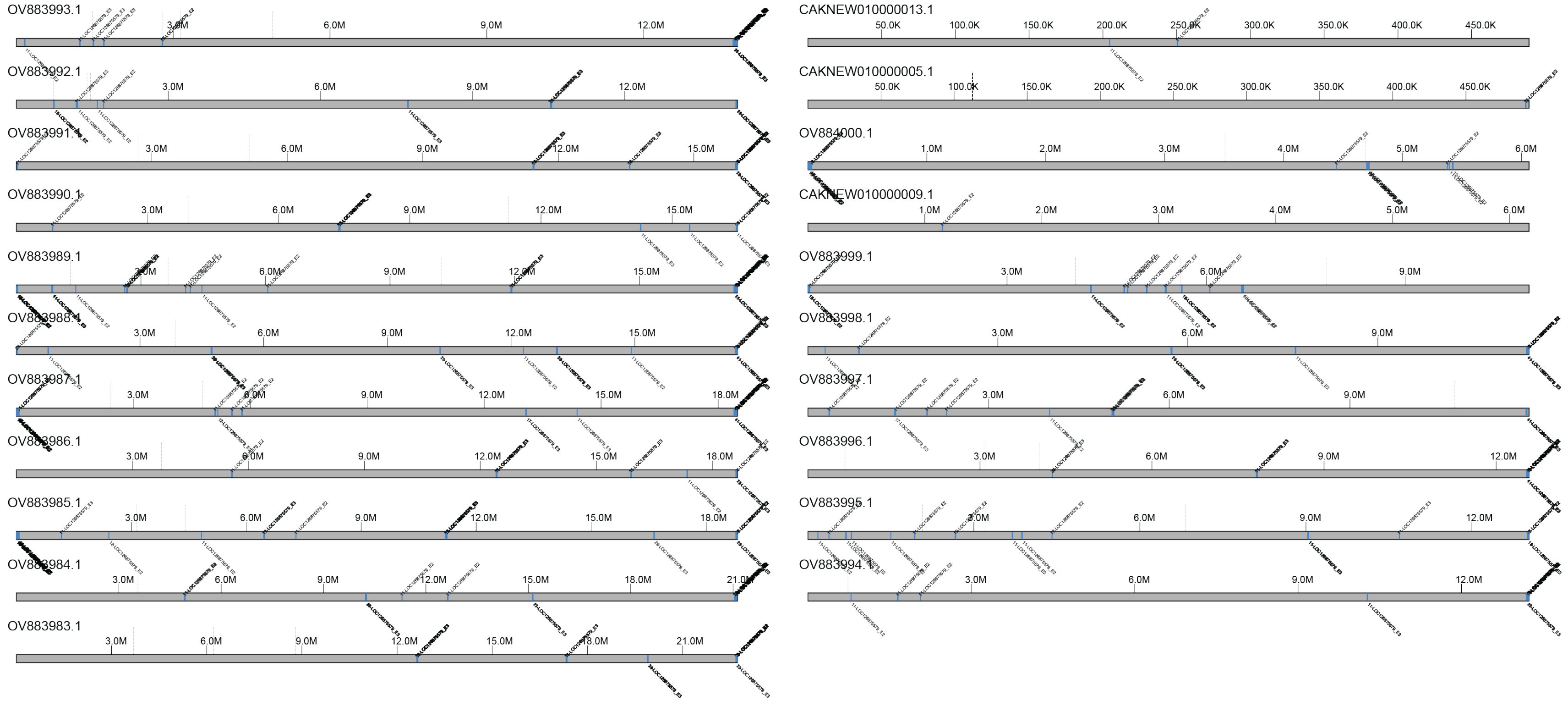


**Figure S17**: Klump plot of LOC126875579 klumps throughout the *Bombus pratorum* reference genome. Blue boxes illustrate LOC126875579 klumps, and dashed lines above each sequence represent gaps in the sequence.

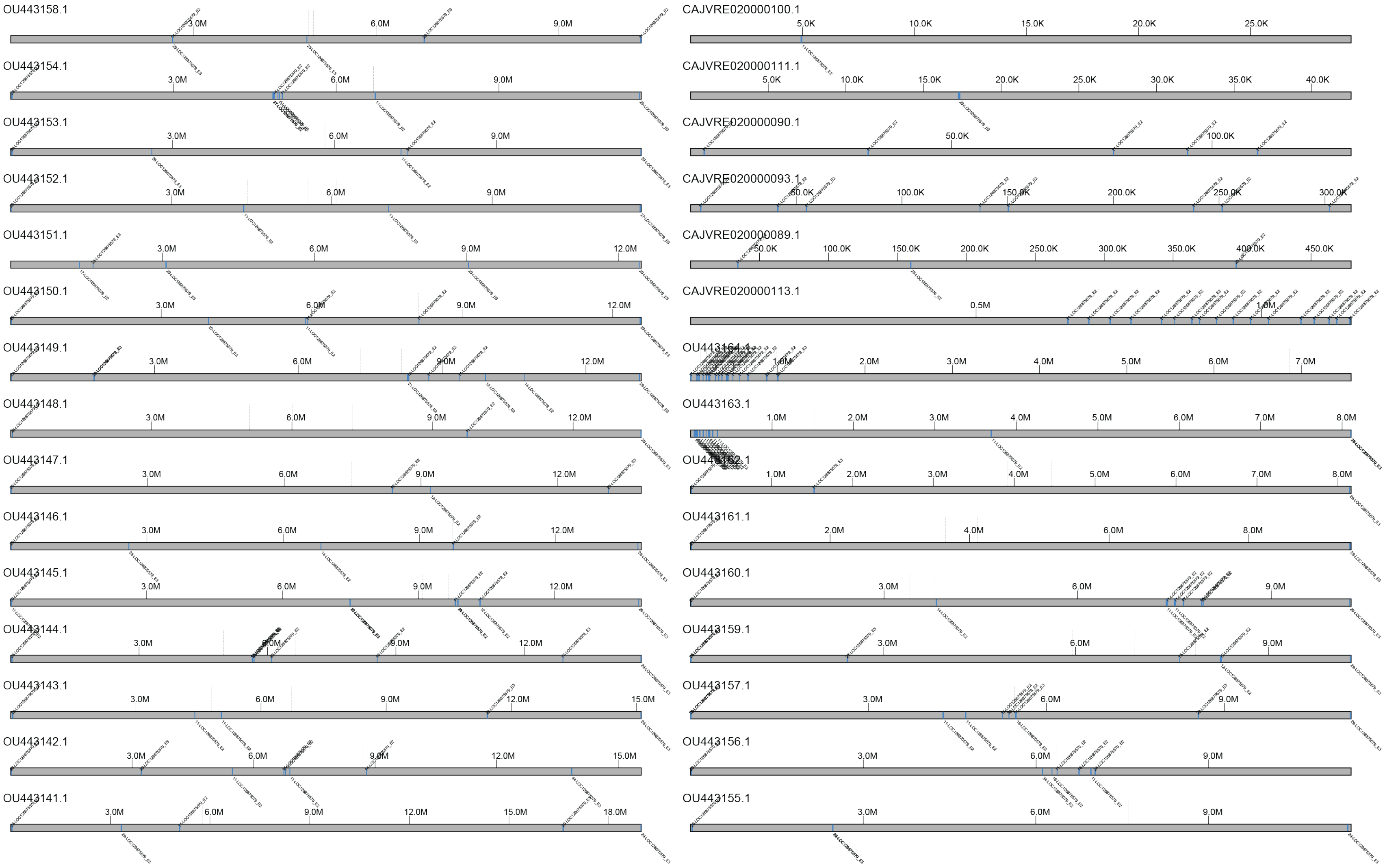


**Figure S18**: Klump plot of LOC126875579 klumps throughout the *Bombus sylvestris* reference genome. Blue boxes illustrate LOC126875579 klumps, and dashed lines above each sequence represent gaps in the sequence.

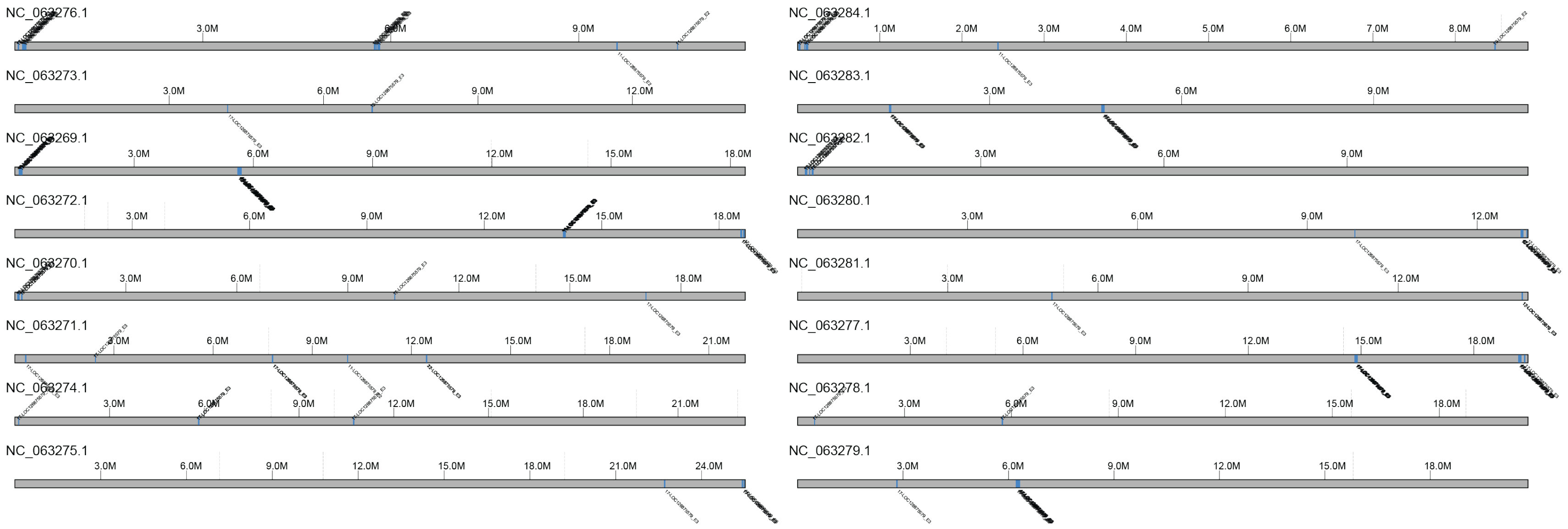


**Figure S19**: Klump plot of LOC126875579 klumps throughout the *Bombus terrestris* reference genome. Blue boxes illustrate LOC126875579 klumps, and dashed lines above each sequence represent gaps in the sequence.

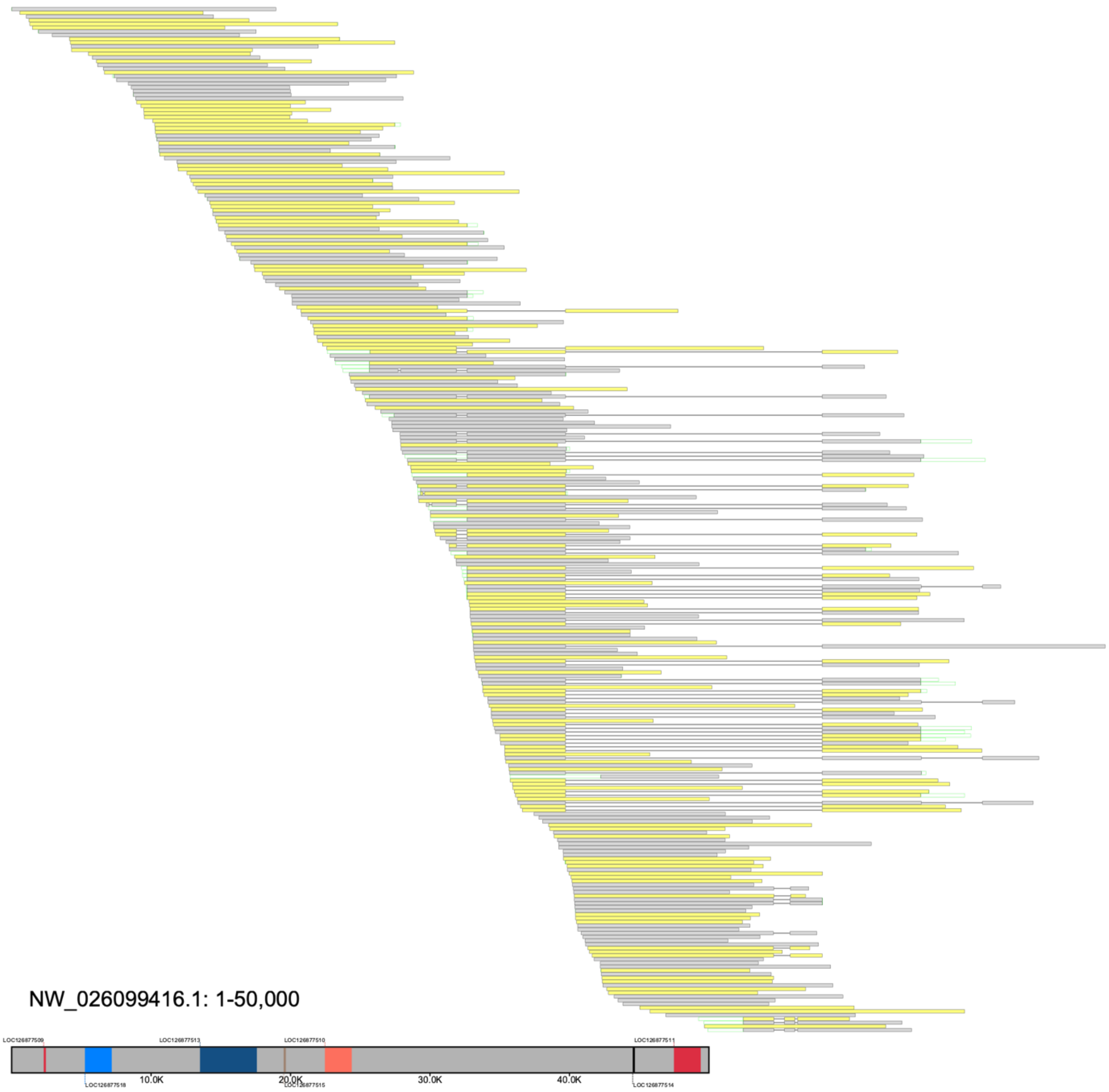


**Figure S20**: A flagged region in the *B. huntii* reference genome after implementing `scan_alignments` using default parameters. Alignments consistent with the reference can be seen at the end of this window where several alignments contain a deletion. We predict that the proportion of overlap between the alignments that are consistent with the reference genome was not sufficient to group them into a single group that is capable of tiling across this region.
